# Functional connectome through the human life span

**DOI:** 10.1101/2023.09.12.557193

**Authors:** Lianglong Sun, Tengda Zhao, Xinyuan Liang, Mingrui Xia, Qiongling Li, Xuhong Liao, Gaolang Gong, Qian Wang, Chenxuan Pang, Qian Yu, Yanchao Bi, Pindong Chen, Rui Chen, Yuan Chen, Taolin Chen, Jingliang Cheng, Yuqi Cheng, Zaixu Cui, Zhengjia Dai, Yao Deng, Yuyin Ding, Qi Dong, Dingna Duan, Jia-Hong Gao, Qiyong Gong, Ying Han, Zaizhu Han, Chu-Chung Huang, Ruiwang Huang, Ran Huo, Lingjiang Li, Ching-Po Lin, Qixiang Lin, Bangshan Liu, Chao Liu, Ningyu Liu, Ying Liu, Yong Liu, Jing Lu, Leilei Ma, Weiwei Men, Shaozheng Qin, Jiang Qiu, Shijun Qiu, Tianmei Si, Shuping Tan, Yanqing Tang, Sha Tao, Dawei Wang, Fei Wang, Jiali Wang, Pan Wang, Xiaoqin Wang, Yanpei Wang, Dongtao Wei, Yankun Wu, Peng Xie, Xiufeng Xu, Yuehua Xu, Zhilei Xu, Liyuan Yang, Huishu Yuan, Zilong Zeng, Haibo Zhang, Xi Zhang, Gai Zhao, Yanting Zheng, Suyu Zhong, Alzheimer’s Disease Neuroimaging Initiative, Cam-CAN, Developing Human Connectome Project, DIDA-MDD Working Group, MCADI, NSPN, Yong He

## Abstract

The lifespan growth of the functional connectome remains unknown. Here, we assemble task-free functional and structural magnetic resonance imaging data from 33,250 individuals aged 32 postmenstrual weeks to 80 years from 132 global sites. We report critical inflection points in the nonlinear growth curves of the global mean and variance of the connectome, peaking in the late fourth and late third decades of life, respectively. After constructing a fine-grained, lifespan-wide suite of system-level brain atlases, we show distinct maturation timelines for functional segregation within different systems. Lifespan growth of regional connectivity is organized along a primary-to-association cortical axis. These connectome-based normative models reveal substantial individual heterogeneities in functional brain networks in patients with autism spectrum disorder, major depressive disorder, and Alzheimer’s disease. These findings elucidate the lifespan evolution of the functional connectome and can serve as a normative reference for quantifying individual variation in development, aging, and neuropsychiatric disorders.

## Introduction

The resting human brain, characterized by intrinsic or spontaneous brain activities, has been increasingly understood from a connectome perspective over the past two decades ^1–5^. The emergence, development, and aging of the intrinsic connectome architecture enables the dynamic reorganization of functional specialization and integration throughout the lifespan, contributing to continuous changes in human cognition and behavior ^6–9^. Understanding the spatiotemporal growth process of the typical functional connectome is critical for elucidating network-level developmental principles in healthy individuals and for pinpointing periods of heightened vulnerability or potential. Disruption of these normative connectome patterns, especially during specific time windows, can predispose individuals to a spectrum of neurodevelopmental ^10–12^, neurodegenerative ^13^, and psychiatric disorders ^14–16^. The growth chart framework provides an invaluable tool for charting normative reference curves in the human brain ^17–20^. Recently, Bethlehem et al. ^18^ delineated the life-cycle growth curves of brain morphometry by aggregating the largest multisite structural magnetic resonance imaging (MRI) dataset to date (101,457 individuals between 115 days post-conception to 100 years of age), marking a significant step toward reproducible and generalizable brain charts. However, the normative growth charts of the functional brain connectome across the human lifespan remain unknown.

Previous studies using task-free functional MRI (fMRI) data have reported age-related characteristics of the functional connectome ^21–23^. However, most of these studies were limited to specific periods of growth with narrow age intervals. For example, data from the perinatal and early postnatal period (e.g., 0-6 years) are rarely included in studies spanning childhood, adolescence, and adulthood; thus missing the opportunity to depict a continuous life-cycle dynamic evolution from gestation to old age. Although a few studies have attempted to include a broader age range from childhood to late adulthood, they have suffered from challenges in robustly estimating normative growth curves due to limited sample sizes (typically < 1,000) ^24–29^. More recently, Rutherford et al. ^30^ have made great strides in establishing a lifespan normative model of the functional connectome using a large sample dataset (∼22,000 individuals aged 2-100 years). However, this work primarily focused on intersystem functional connectivity using population-based system-level atlas. Furthermore, there are large inconsistencies in the literature regarding functional connectivity trajectories, with no consensus emerging on the developmental directions and growth milestones. In particular, Cao et al. ^25^ reported that global functional connectivity in the whole brain peaks at around 30 years of age, whereas other studies suggest earlier peaks ^24^ or show a continuous decline across the lifespan ^31^. Different trends have been observed for sensorimotor regions, with reports of ascending ^32^, descending ^33^, and stable ^34^ developmental trajectories from childhood to adolescence. Similarly, connectivity patterns between the default and frontoparietal networks have been reported to both increase ^35^ and decrease ^36, 37^ during this period. Such discrepancies between studies are likely due to the high sensitivity of high-dimensional fMRI data to variations in scanner platforms and sequences, image quality, data processing, and statistical methods, as well as the population heterogeneity of cohorts ^6^. This underscores the paramount importance of large sample sizes, rigorous data quality control procedures, consistent data processing protocols, and standardized statistical modeling frameworks to accurately characterize growth curves of the functional connectome across the lifespan.

To address this gap, we assembled a large multimodal neuroimaging dataset with rigorous quality control, consisting of cross-sectional task-free fMRI and structural MRI data from 33,250 individuals ranging in age from 32 postmenstrual weeks to 80 years, collected from 132 global sites (Fig. 1a). We conducted a comprehensive network modeling analysis to delineate the nonlinear growth patterns of the functional connectome across multiple scales. We began by characterizing lifespan growth in the overall patterns of the global functional connectome, revealing important life-course milestones. We then constructed continuous age-related, system-level atlases across the lifespan and further provided a previously unreported portrayal of the distinct growth patterns across brain systems. Next, we sought to elucidate the spatiotemporal principles governing connectome growth at a finer regional scale. Finally, we investigated the potential clinical value of the established connectome-based normative models. We selected autism spectrum disorder (ASD, N = 414), major depressive disorder (MDD, N = 622), and Alzheimer’s disease (AD, N = 180) as representative conditions characterized by network dysfunction across different life stages. These conditions typically manifest in childhood, adolescence/adulthood, and older adulthood, respectively ^16, 38–40^. Using individual deviation scores relative to the 50th percentile, we presented a multiscale characterization to quantify the individual heterogeneity of patients with ASD, MDD, or AD.

**Fig. 1.**
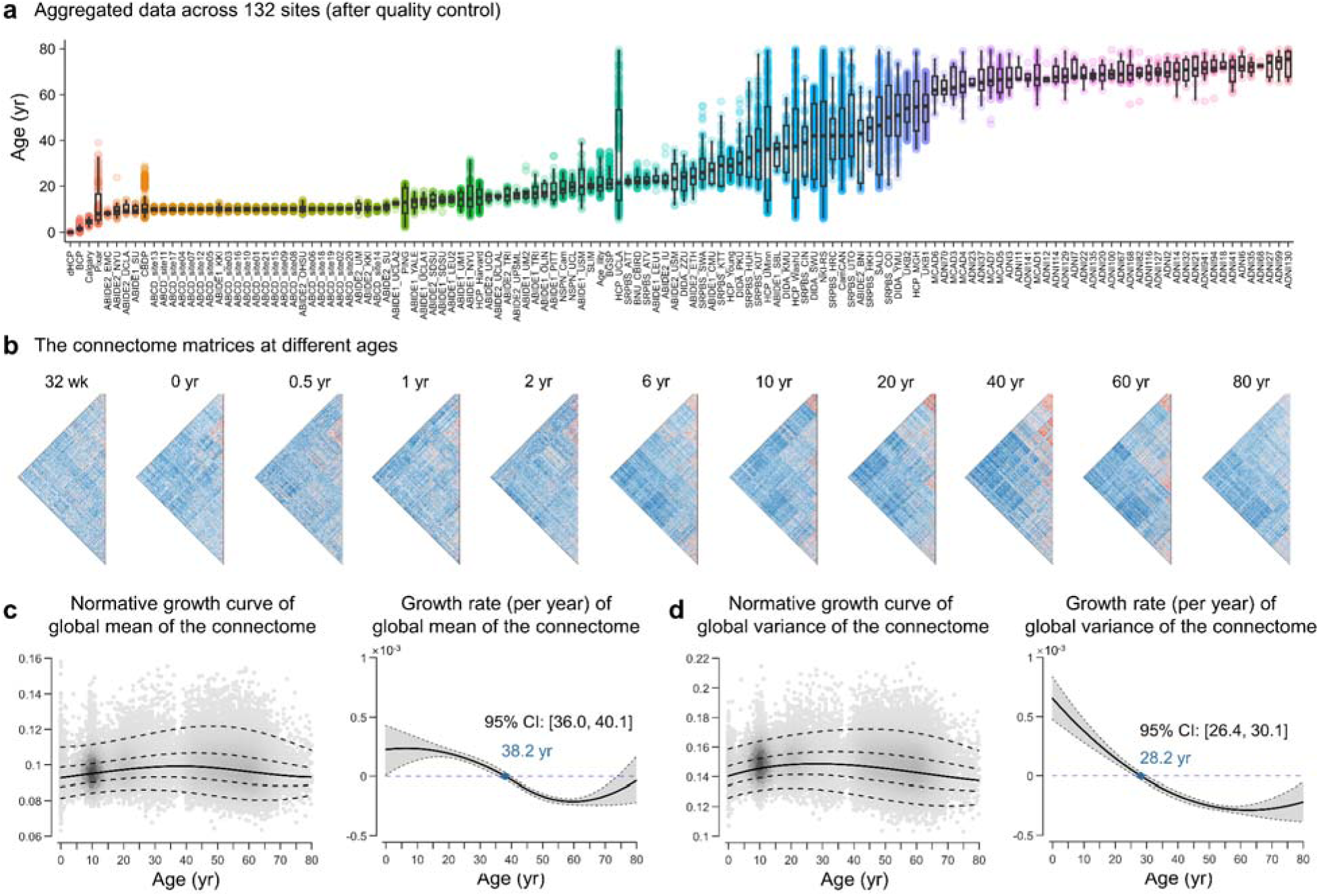
Normative growth patterns of the functional connectome at a global level over the lifespan. **a**, Quality-controlled MRI data from 132 scanning sites comprising 33,250 healthy participants who collectively spanned the age range from 32 postmenstrual weeks to 80 years. Box plots show the age distribution of participants at each site of data acquisition. The detailed participant demographics and acquisition parameters of each site are provided in Supplementary Tables 1 and 2, respectively. **b**, The functional connectome matrices of representative participants at different ages. **c**, Normative growth curve (left panel) and growth rate (right panel) of the global mean of the connectome as estimated by GAMLSS. The median (50th) centile is represented by a solid line, while the 5th, 25th, 75th, and 95th centiles are indicated by dotted lines. The growth rate is characterized by the first derivative of the median centile line. The gray shaded areas represent the 95% confidence interval, which was estimated by bootstrapping 1,000 times (see Methods for details). **d**, Normative growth curve (left panel) and growth rate (right panel) of global variance of the connectome. wk, week; yr, year.

## Results

We initially aggregated 44,030 participants with multimodal structural MRI and task-free fMRI data. After a rigorous quality control process (for details, see the Methods and Supplementary Figs 1 and 2), we obtained a final sample of 34,466 participants with high-quality imaging data, including 33,250 healthy individuals (Fig. 1a) and 1,216 patients. The detailed demographics and acquisition parameters of the datasets are provided in Supplementary Tables 1 and 2, respectively. Using the standardized and highly uniform processing pipeline (Methods and Supplementary Fig. 3), we obtained the surface-based preprocessed blood oxygenation level-dependent (BOLD) signals in fsaverage4 space for each participant (4,609 vertices in total). We then constructed a vertexwise 4,609×4,609 functional connectome matrix by calculating Pearson’s correlation coefficient between the time courses of each vertex. Figure 1b shows the functional connectome matrices of representative participants at different ages. Next, we examined the individual connectome at the global, system, and vertex levels. In accordance with the recommendations of the World Health Organization recommendation ^41^, the age-related nonlinear growth patterns were described using the generalized additive model for location, scale, and shape (GAMLSS) ^41, 42^, based on cross-sectional data from healthy populations (N = 33,250). Sex and in-scanner head motion (mean framewise displacement) were included as fixed effect covariates, and the scanner site was included as a random effect covariate. GAMLSS provides a robust framework for modeling nonlinear growth curves and has been widely used in neurodevelopmental studies ^18, 43–45^. To assess the rate of growth (velocity) and inflection points, we calculated the first derivatives of the lifespan growth curves. The GAMLSS specifications, model estimations, and model evaluations are detailed in the Methods section.

### Mapping the normative growth of the global functional connectome across the lifespan

To provide basic developmental and aging insights into the global functional connectome, we first characterized the normative growth patterns of the global mean and variance (estimated by standard deviation) of the functional connectome. The lifespan curve of the global mean of functional connectome (Fig. 1c) exhibited a nonlinear increase from 32 postmenstrual weeks onward, peaking in the late fourth decade of life (38.2 years, 95% bootstrap confidence interval (CI) 36.0-40.1), followed by a nonlinear decline. This growth curve is primarily driven by age-related changes of middle- and long-range connections (Supplementary Fig. 4). The global variance of functional connectome (Fig. 1d) also exhibited a nonlinear growth pattern, reaching its peak in the late third decade of life (28.2 years, 95% bootstrap CI 26.4-30.1). The utilization of the GAMLSS enabled the delineation of normative growth curves for interindividual variability ^18^ in the two global measures (Supplementary Result 1 and Supplementary Fig. 5a).

The curves demonstrated a slight decline in inter-individual variability during the initial stages of early development, a gradual increase until the late sixth decade of life (peaking at 55.2 years, 95% CI [53.9, 56.0] for the global mean; peaking at 56.8 years, 95% CI [55.1, 58.1] for the global variance), and then a rapid decline. These nonlinear growth patterns in the global connectome measures indicated a temporally coordinated manner across the lifespan.

### Lifespan growth of system-specific organization in the functional connectome

Functional segregation and integration are two fundamental organizational principles of the human brain connectome ^1^. To understand the lifespan growth patterns of functional segregation and integration, we established the normative models of the functional connectome at the systems level. The first step was to parcellate the cortex into distinct functional systems for each participant. Converging evidence has shown that relying on population-level atlases for individual analysis overlooks crucial intersubject variability in functional topography organization ^46–49^. This oversight leads to the misinterpretation of spatial distribution differences as system-level disparities ^47, 50^, thereby increasing the risk of inaccuracies in mapping both intra- and intersystem connectivity. Moreover, although previous studies of fetal and infant brains have elucidated the early emergence of basic forms of large-scale functional systems, including the visual ^51–54^, somatomotor ^51–54^, dorsal attention ^55, 56^, ventral attention ^51^, frontoparietal ^52, 54, 56^, and default mode networks ^51–54, 56^, the functional architecture of an individual’s system undergoes dramatic refinement and reorganization over the protracted life course ^21, 57^. To increase the precision of the construction of individual-specific functional networks, it is essential to establish a set of continuous growth atlases with accurate system correspondences across the life course.

To address this issue, we proposed a Gaussian-weighted iterative age-specific group atlas (GIAGA) generation approach (see Methods and Supplementary Fig. 6a). The iterative refinement process is central to this approach. Briefly, we first divided all participants aged 32 postmenstrual weeks to 80 years into 26 distinct age groups. Yeo’s adult atlas ^58^ was then used as a prior to generate a personalized parcellation for each participant in a given age group. These personalized parcellations were further aggregated to construct an age-specific population-level atlas, where the contribution of participants was weighted according to their age position within a Gaussian probability distribution. This process was repeated until the age-specific population-level atlas converged, resulting in a set of age-specific brain atlases across the lifespan (Fig. 2a, Supplementary Figs 7 and 8). Validation analysis revealed greater global homogeneity when using these age-specific group atlases than using the adult-based group atlas across all age groups (all p < 10^-9^, Bonferroni-corrected, Supplementary Fig. 9), particularly evident during early development. Notably, each of the 26 brain atlases was parcellated into seven canonical functional networks. For each network, we calculated the network size ratio, measured by the proportion of vertices, and the distribution score, defined by the number of spatially discontinuous subregions (Fig. 2b). We found that the default mode (DM), frontoparietal (FP), and ventral attention (VA) networks showed a slight expansion in network size during the first month of life, while their distribution scores developed until early childhood (4-6 years). In contrast, the somatomotor (SM), visual (VIS), and dorsal attention (DA) networks showed a relatively stable pattern of network size and network discretization throughout the lifespan. A hierarchical clustering analysis of these system-level brain atlases revealed three overarching patterns. Cluster I covered atlases from 34 postmenstrual weeks to 1 month, cluster II covered atlases from 3 months to 24 months, and cluster III covered atlases from 4 years to 80 years of age (Supplementary Fig. 10). To further quantify the growth patterns of the whole-cortical atlas and the system-specific atlases, we computed their network similarity to the designated reference atlas using both the overlay index and the Dice coefficient (Methods). The reference atlas was derived from the average of eight adult-like atlases, identified as a homogeneous cluster of 18- to 80-year-old atlases (Supplementary Fig. 10). We found that the overall similarity of the whole-cortical atlas exhibited a rapid increase during the first two decades of life, followed by a plateau, and a subsequent slight decrease with age (Fig. 2c). At the system level, we observed that both the VIS and SM networks exhibited adult-like patterns (80% similarity) in the perinatal period, whereas the DM, FP, DA, and VA networks developed adult-like patterns (80% similarity) at 4-6 years of age (Fig. 2d and 2e).

**Fig. 2.**
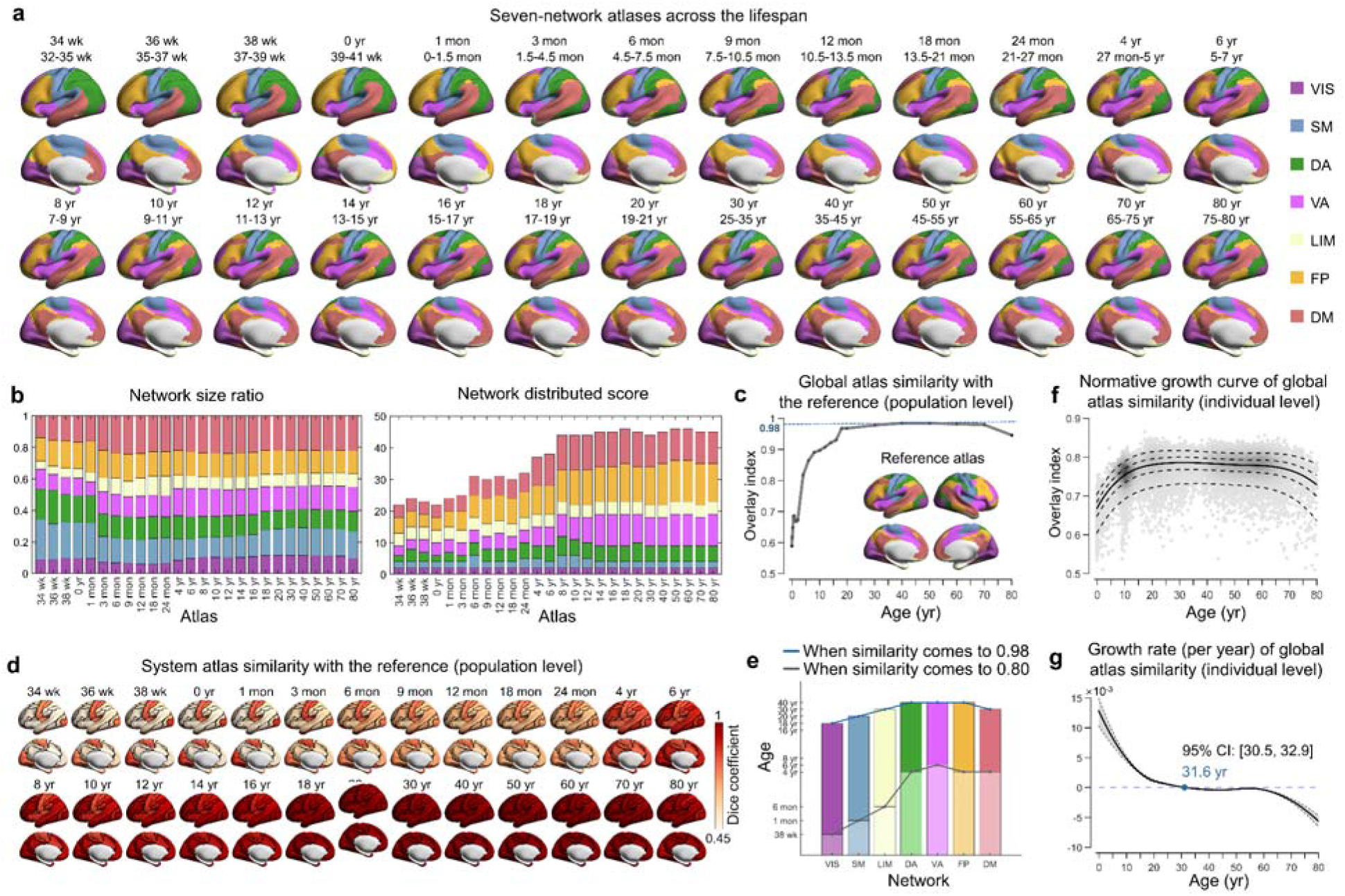
Population-level and individual-level functional atlases throughout the lifespan. **a**, Employing the Gaussian-weighted iterative group atlas generation approach (for details, see Methods and Supplementary Fig. 6a), the lifespan set of seven-network functional atlases from 32 postmenstrual weeks to 80 years was established (26 atlases in total). Only the left hemisphere is displayed here; for the whole-cortical atlases, refer to Supplementary Figs 7 and 8. Labels of each system were mapped onto the HCP fs_LR_32k surface and visualized using BrainNet Viewer ^59^. **b**, Network size ratio and network distribution score of each system in all age-specific group atlases. The network size ratio was calculated as the vertex number of the system divided by the total cortical vertex number. The network distribution score was measured by the number of spatially discontinuous subregions (≥ 5 vertices) in the system. **c**, Global similarity of each age-specific group atlas with the reference atlas across the lifespan. The degree of global similarity was defined as the number of vertices with the same label in the two atlases divided by the total number of vertices in both atlases. **d**, System similarity of each age-specific group atlas with the corresponding system in the reference atlas across the lifespan. System similarity was quantified using the Dice coefficient. **e**, The ages at which the system similarity of each age-specific group atlas reached 0.8 and 0.98. **f-g**, Normative growth curve and growth rate of global atlas similarity with the reference atlas when using personalized functional atlas for each participant. The gray shaded areas represent the 95% confidence interval, which was estimated by bootstrapping 1,000 times. VIS, visual; SM, somatomotor; DA, dorsal attention; VA, ventral attention; LIM, limbic; FP, frontoparietal; DM, default mode. wk, week; mon, month; yr, year.

Based on the age-specific group atlases established above, we proceeded to map individual-level functional systems for each participant. Specifically, we used an iterative parcellation procedure (see Methods and Supplementary Fig. 6b), as proposed by Wang et al. ^60^, which has been demonstrated to accurately identify personalized functional networks in both healthy ^47, 60^ and diseased individuals ^61–63^. As expected, the individual-level atlases exhibited significantly greater global homogeneity than both the age-specific group atlases (all p < 10^-9^, Bonferroni-corrected) and the adult-based group atlas (all p < 10^-8^, Bonferroni-corrected), regardless of the age groups considered (Supplementary Fig. 9). Consistent with the growth pattern observed in the age-specific group atlas (Fig. 2c), the global similarity of the individualized atlas to the reference increased from 32 postmenstrual weeks and reached a peak in adulthood (31.6 years, 95% bootstrap CI 30.5-32.9) (Fig. 2f and 2g).

Using the person-specific network mapping approach, which integrates individual-level iterative processes with the age-specific group atlases, we characterized the lifespan growth patterns of within-system connectivity (functional segregation) and between-system connectivity (functional integration) (Supplementary Result 2, Supplementary Figs 11 and 12). To further quantify the differences in within-system connectivity relative to between-system connectivity, we calculated the system segregation index for each brain system ^64^. This index measures the difference between mean within-system connectivity and mean between-system connectivity as a proportion of mean within-system connectivity ^64^ (Methods). Interestingly, global segregation across all systems peaked in the third decade of life (25.7 years, 95% bootstrap CI 24.8-26.8) (Fig. 3a). At the system level, different networks manifested distinct nonlinear growth patterns (Fig. 3b-3d). The primary VIS network consistently showed the greatest segregation across all ages (Fig. 3b and 3c), suggesting that the VIS network is more functionally specialized and relatively less integrated in inter-network communication compared to other systems. The DA and VIS networks exhibited similar trends in life-cycle growth patterns, peaking in early childhood and pre-adolescence, respectively (Fig. 3b and 3c). The DM and FP networks showed the lowest levels of segregation in the early stages of neurodevelopment (Fig. 3b and 3c). However, segregation increased rapidly with age peaks at the end of the third decade and decreased rapidly in the late stages of senescence (Fig. 3b-3d). Finally, the SM and VA networks showed similar growth patterns of system segregation, increasing and decreasing moderately over the lifetime (Fig. 3b-3d).

**Fig. 3.**
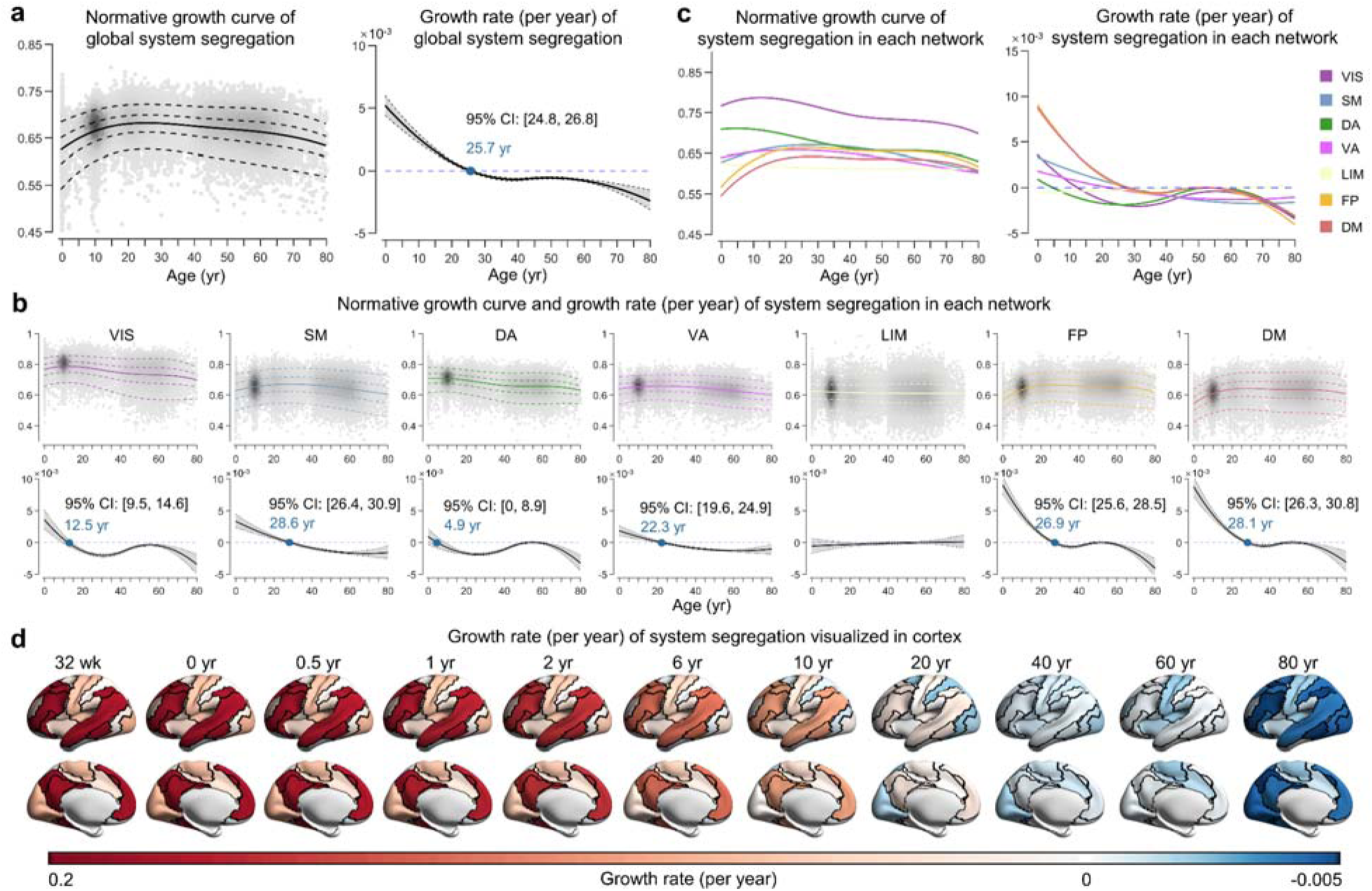
Lifespan normative growth patterns of brain system segregation. **a**, Normative growth curve and growth rate of global system segregation. The peak occurred in the third decade of life (25.7 years, 95% bootstrap confidence interval 24.8-26.8). The gray shaded areas represent the 95% confidence interval, which was estimated by bootstrapping 1,000 times. **b-c**, Normative growth curves and growth rate of system segregation for each network. The median (50th) centile is represented by a solid line, while the 5th, 25th, 75th, and 95th centiles are indicated by dotted lines. The key inflection points are marked in blue font. **d**, Growth rate of system-specific segregation visualized in the cortex, with black lines depicting system boundaries. The values of each system are mapped and visualized on the HCP fs_LR_32k surface. VIS, visual; SM, somatomotor; DA, dorsal attention; VA, ventral attention; LIM, limbic; FP, frontoparietal; DM, default mode. wk, week; yr, year.

### Lifespan growth of functional connectivity at the regional level reveals a spatial gradient pattern

Having identified distinct growth patterns in different brain systems, we further explored the more nuanced spatiotemporal growth patterns of the functional connectome at the regional level. First, we plotted the normative growth curves of each vertex’s functional connectivity strength (FCS) by calculating the average connectivity with all other vertices. Figure 4a shows the curves for several vertices located in different brain regions, and Figure 4b shows the fitted FCS and its growth rate across the cortex. Notably, the most pronounced changes in functional connectivity at the regional level occurred within the first decade of life. We then sought to elucidate how the overall growth patterns varied spatially across the cortex by mapping the primary spatial axis of FCS development. To this end, we used a principal component analysis (PCA) on the zero-centered 50th centiles of the growth curves. The first PC, accounting for 60.4% of the variance, was identified as the dominant axis of regional functional connectivity growth (Fig. 4c). This axis captured a hierarchical spatial transition, starting from primary sensorimotor and visual cortices and culminating in higher-order association regions, including the angular gyrus, precuneus, temporal, and prefrontal cortices. To better illustrate the spatiotemporal pattern of growth curves throughout the cortex, we segmented the main growth axis into 20 equal bins and averaged the curves for vertices within each bin. A continuous spectrum of curves along the lifespan axis is shown in Fig. 4d.

**Fig. 4.**
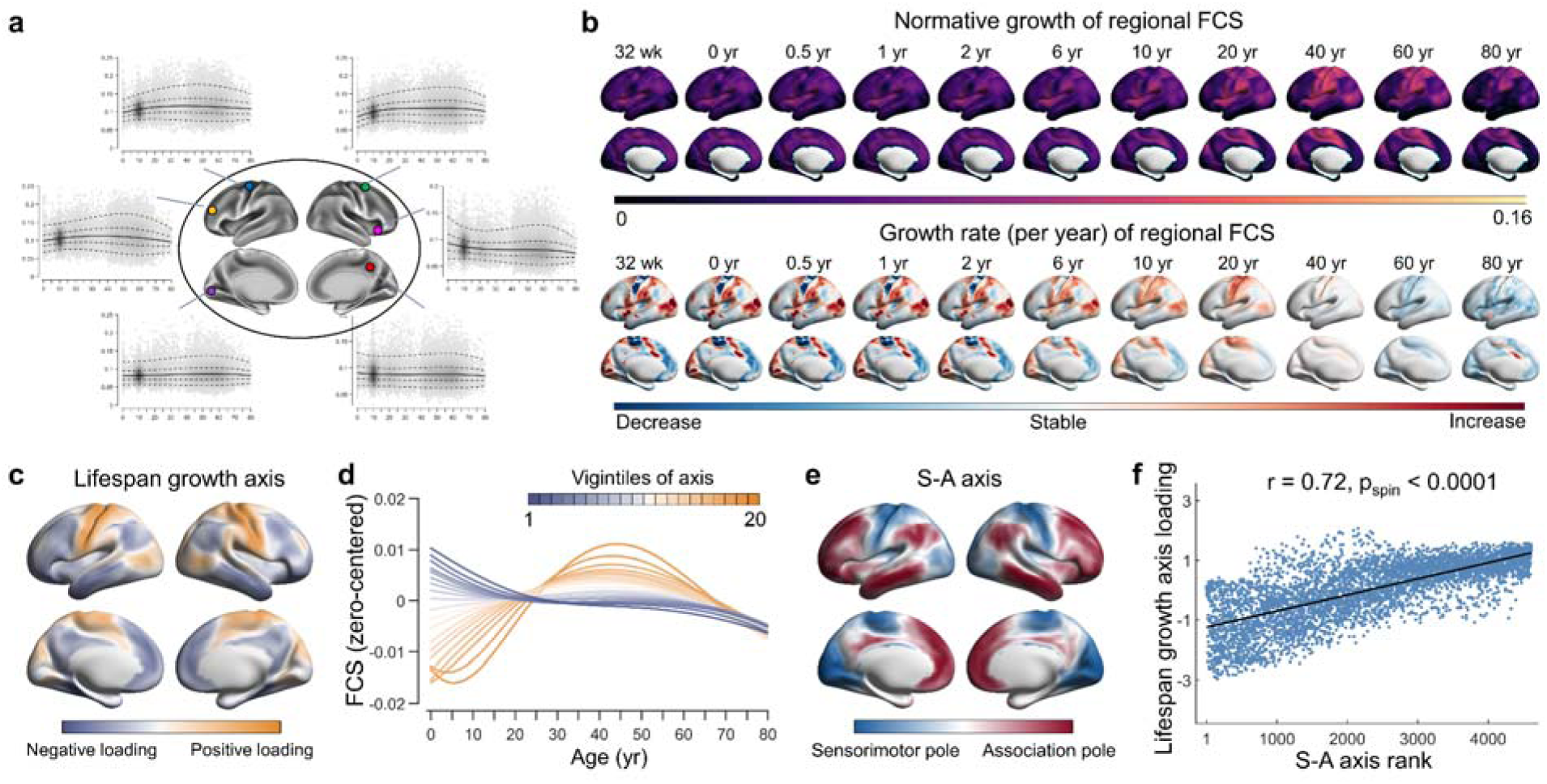
Lifespan normative growth patterns of regional functional connectivity strength. **a**, Normative growth curves of example vertices from different regions. **b**, The fitted 50th centiles (top panel) and their growth rates (bottom panel) for all vertices at representative ages. **c**, The lifespan growth axis of brain functional connectivity, represented by the first principal component from a PCA on regional level FCS curves. **d**, Based on the lifespan principal axis, all vertices across the brain were equally divided into 20 bins. The zero-centered curves of all vertices within each bin were averaged. The first vigintile (depicted in darkest yellow) represents one pole of the axis, while the twentieth vigintile represents the opposite pole (depicted in darkest blue). The patterns of growth curves vary continuously along the axis, with the greatest differences observed between the two poles. **e,** The sensorimotor-association (S-A) axis, as formulated by Sydnor et al. ^66^, represents a cortical continuum that transitions from primary regions to transmodal areas. **f,** A strong correlation was observed between the lifespan principal growth axis and the S-A axis (r = 0.72, p_spin_ < 0.0001) (linear association shown with 95% confidence interval). All brain maps were mapped to the HCP fs_LR_32k surface for visualization. FCS, functional connectivity strength; wk, week; yr, year.

The cortical landscape of the human brain is organized by a fundamental gradient known as the sensorimotor-association (S-A) axis ^65^. This axis spans from primary cortices critical for sensory and motor functions to advanced transmodal regions responsible for complex cognitive and socioemotional tasks. It has been shown to play an important role in shaping neurodevelopmental processes ^66–68^. Here, we sought to investigate the extent to which our defined growth axis aligns with the classic S-A axis as formulated by Sydnor et al. ^66^. (Fig. 4e). Using a spin-based spatial permutation test ^69^, we found a significant association between the main growth axis and the S-A axis (r = 0.72, p_spin_ < 0.0001) (Fig. 4f). This finding suggests that the spatiotemporal growth of the functional connectome throughout the human lifespan follows the canonical sensorimotor-association organization.

### Sex differences in lifespan growth patterns

It is becoming increasingly evident that sex differences exert a significant influence on brain development and aging ^70, 71^. In GAMLSS modeling, we included a sex effect as an additional variable to establish lifespan normative growth curves. We characterized the sex-stratified growth curves and interindividual variability curves of the functional connectome (Supplementary Result 3, Supplementary Figs 13 and 14). Specifically, we observed that the global mean of the functional connectome was significantly greater in males than in females (p_FDR_ = 0.0002), thereby confirming and extending conclusions from previous studies ^72, 73^.

Conversely, the global variance of the connectome was greater in females than in males (p_FDR_ = 0.0009). Furthermore, females showed greater global system segregation (p_FDR_ = 10^-24^) and system-specific segregation in the VIS, VA, FP, and DM networks (all p_FDR_ < 0.01), but lower system-specific segregation in the SM and limbic (LIM) networks (all p_FDR_ < 10^-32^) than males. At the regional level, the lateral and medial parietal cortex and lateral prefrontal cortex showed greater FCS in females, whereas the sensorimotor cortex, medial prefrontal cortex, and superior temporal gyrus showed greater FCS in males (p_FDR_ < 0.05). These results are compatible with a previous study employing seed-based and independent component analysis (ICA)-based functional connectivity analysis ^23^. Additionally, in a recent study, Zhang et al. ^74^ used a large data set (36,531 participants from the UK Biobank, mean age 69) to report that females had lower functional connectivity in somatosensory/premotor regions and greater functional connectivity in the inferior parietal and posterior cingulate cortex, which aligns with our findings. The detailed statistical values of the sex variable within each normative model are presented in Supplementary Tables 3 and 4. The sex differences in the interindividual variability curves are detailed in the Supplementary Result 3.

### Identifying individual heterogeneity in brain disorders using connectome-based normative models

Recent studies have highlighted the potential of normative models to disentangle the inherent heterogeneity in clinical cohorts by enabling statistical inference at the individual level ^18, 75–81^. This approach enables the quantification of individual deviations of brain phenotypes from normative expectations, thereby providing unique insights into the typicality or atypicality of individual’s brain structure or function. To validate the clinical value of our connectome-based normative models, we selected three representative brain disorders characterized by connectome dysfunction, each manifesting at distinct life stages. These were ASD, which mainly presents in early development; MDD, which mainly presents in adolescence and adulthood; and AD, which mainly presents in older adulthood.

We characterized the individual deviation z-scores (age- and sex-specific) of the functional metrics at the global, system, and regional levels in patients with ASD (N_ASD_ = 414, aged 5-59 years), MDD (N_MDD_ = 622, aged 11-77 years), or AD (N_AD_ = 180, aged 51-80 years), and their matched healthy controls (HCs). The standard protocol for normative modeling ^82^ emphasizes the importance of incorporating some control samples from the same imaging sites as the patients into the testing set. This approach verifies that the observed case□control differences are not due to the analysis with controls in the training set and cases in the testing set ^78, 82^. This approach also allows for the estimation of site effects within the case□control datasets. In the present study, we reconstructed the connectome-based normative models for all three disorders using the same set of healthy participants. Specifically, we randomly divided the HCs of all case□control datasets (N_HC_ = 591 in ASD datasets, N_HC_ = 535 in MDD datasets, and N_HC_ = 187 in AD datasets) in half, stratified by age, sex, and site. The training set (N = 32,591), which was used to construct the normative model, consisted of half of the HCs (N_train_ = 654) and all samples from other datasets (N = 31,937). The testing set, comprising remaining half of the HCs (N_test_ = 659) and the patient cases, was used as a completely independent set to determine their deviation scores. This process was repeated 100 times, generating 100 new normative models and 100 sets of deviation scores. We observed a high degree of stability of both normative curves and patient’s deviation scores across the 100 repetitions (average r > 0.95 and average mean square error [MSE] < 0.2 for all functional metrics, see Supplementary Fig. 15, Supplementary Tables 5 and 6). We then averaged the 100 sets of deviation scores for patients in each disease group, and then assessed the extreme deviations (z > |2.6|) for each metric. Among the ASD patients, 92% had at least one metric with an extreme negative deviation, and 32% had at least one metric with an extreme positive deviation (Supplementary Fig. 16). For MDD patients, the percentages were 89% and 39%, respectively, and for the AD patients, they were 61% and 25%, respectively. Furthermore, we calculated the proportion of patients with extreme deviations in each metric and found that no more than 10% of the patients had extreme deviations in any single metric (Fig. 5a, Supplementary Fig. 16). These results highlight the considerable individual heterogeneity within each disease group.

**Fig. 5.**
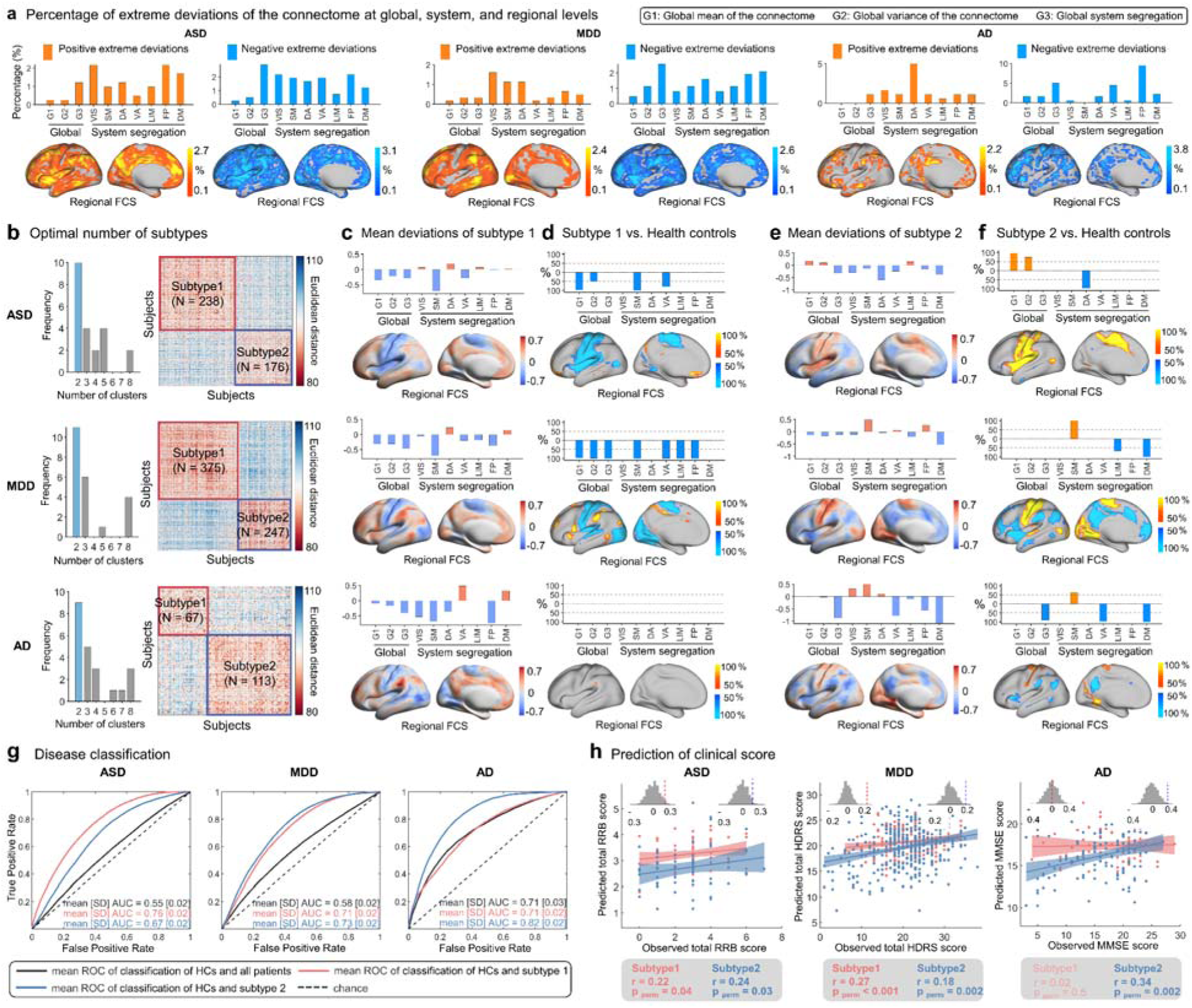
Clinical relevance of connectome-based deviation patterns in three brain disorders. **a**, Percentage of patients with extreme deviations. Subplots from left to right display the percentage of patients with extreme positive and negative deviations in ASD, MDD, and AD. The bar plot shows the percentage in global mean of the connectome (G1), global variance of the connectome (G2), global system segregation (G3), and system-specific segregation. The brain map shows the percentage of regional-level FCS. OrangeDyellow represents extreme positive deviations, while blue represents extreme negative deviations. **b**, The optimal number of subtypes (left panel) and the similarity matrix of deviation patterns across patients (right panel) for each disorder. **c**, Mean deviation patterns in patients in subtype 1 of each disorder. **d**, Individual deviation scores of patients in subtype 1 were compared to the median of healthy controls (HCs). For each metric, the significance of the median differences between the case group and HCs was assessed using the MannDWhitney U test. *P*-values were adjusted for multiple comparisons using FDR correction across all possible pairwise tests (p < 0.05). The color bar represents the proportion of tests that passed the significance threshold in 100 comparisons. **e**, Mean deviations pattern in patients in subtype 2 of each disorder. **f**, Individual deviation scores of patients in subtype 2 were compared to the median of HCs. **g**, Disease classification performance based on individual deviation patterns using support vector machine analysis. **h**, Prediction accuracy of clinical scores based on individual deviation patterns using support vector regression analysis. All brain maps were mapped to the HCP fs_LR_32k surface and are shown in the left hemisphere. For whole-cortex visualizations, refer to Supplementary Figs 16-18. VIS, visual; SM, somatomotor; DA, dorsal attention; VA, ventral attention; LIM, limbic; FP, frontoparietal; DM, default mode. ROC, receiver operating characteristic; AUC, area under the curve; RRB, Repetitive Restrictive Behavior; HDRS, Hamilton Depression Rating Scale; MMSE, Mini-Mental State Examination.

Using the k-means clustering algorithm, we identified two subtypes for ASD (N_ASD1_ = 238 for subtype 1, N_ASD2_ = 176 for subtype 2), two for MDD (N_MDD1_ = 375 for subtype 1, N_MDD2_ = 247 for subtype 2), and two for AD (N_AD1_ = 67 for subtype 1, N_AD1_ = 113 for subtype 2) (Fig. 5b). For each disorder, different subtypes showed distinct patterns of deviation and case□control differences in the functional connectome (Fig. 5c-f, Supplementary Figs 17 and 18). Specifically, ASD subtype 1 showed greater positive deviations in the bilateral ventral prefrontal cortex and negative deviations primarily in the sensorimotor and insular cortices in comparison with HCs (p_FDR_ < 0.05, Fig. 5c-d). In contrast, ASD subtype 2 exhibited greater positive deviations in the sensorimotor and insular cortices (p_FDR_ < 0.05, Fig. 5e-f). For MDD, subtype 1 patients showed greater positive deviations in the lateral frontal and parietal regions, and insular cortices, and greater negative deviations in the visual and sensorimotor cortices (p_FDR_ < 0.05, Fig. 5c-d). MDD subtype 2 patients showed greater positive deviations in the visual and sensorimotor cortices, and greater negative deviations in the lateral and medial prefrontal and parietal regions, and the insula (p_FDR_ < 0.05, Fig. 5e-f). For AD, subtype 1 showed very few positive or negative deviations (Fig. 5c-d), but subtype 2 showed greater positive deviations in the visual and sensorimotor cortices, and negative deviations in the lateral and medial parietal regions and the insula (p_FDR_ < 0.05, Fig. 5e-f).

We further investigated the classification performance of each disorder with and without subtyping, characterized by the mean area under the curves (AUCs) (Fig. 5g, Supplementary Fig. 19). Specifically, the mean AUCs for ASD subtypes 1 and 2 were 0.76 and 0.67, respectively (all p_FDR_ < 0.001, permutation tests), but 0.55 without subtyping (p_FDR_ < 0.05, permutation tests). The mean AUCs for MDD subtypes 1 and 2 were 0.71 and 0.73, respectively (all p_FDR_ < 0.001, permutation tests), but 0.58 without subtyping (p_FDR_ < 0.05, permutation tests). The mean AUCs for AD subtypes 1 and 2 were 0.71 (p_FDR_ < 0.05, permutation tests) and 0.82 (p_FDR_ < 0.001, permutation tests), respectively, but 0.71 without subtyping (p_FDR_ < 0.001, permutation tests).

Furthermore, we investigated the potential of the connectome deviations to predict clinical scores (Fig. 5h, Supplementary Fig. 20). For ASD, the patterns of connectome-based deviations in patients predicted total Repetitive Restrictive Behavior (RRB) scores for each subtype (r = 0.22, p_perm_ = 0.04 for subtype 1; r = 0.24, p_perm_ = 0.03 for subtype 2). However, the prediction was not significant without ASD subtyping (r = 0.04, p_perm_ = 0.3). For MDD, the connectome-based deviation patterns showed significant predictive accuracy for total Hamilton Depression Rating Scale (HDRS) score, both with and without subtyping (r = 0.19, p_perm_ < 0.001 for all patients; r = 0.27, p_perm_ < 0.001 for subtype 1; r = 0.18, p_perm_ = 0.002 for subtype 2). For AD, the prediction of the Mini-Mental State Examination (MMSE) score was significant without subtyping (r = 0.33, p_perm_ < 0.001), but only for AD subtype 2 (r = 0.02, p_perm_ = 0.5 for subtype 1; r = 0.34, p_perm_ = 0.002 for subtype 2). These results demonstrate the clinical relevance of connectome-based normative modeling in identifying disease subtypes, as evidenced by enhanced performance in both disease classification and prediction of symptomatic scores.

### Sensitivity analyses

The lifespan growth patterns of functional connectomes were validated at the global, system, and regional levels using various analysis strategies (for details, see Methods). Each validation strategy yielded growth patterns that highly matched the main results (Supplementary Tables 7-12 and Supplementary Fig. 21). (*i*) To validate the potential effects of head motion, the analyses were reperformed using data from 24,494 participants with a stricter quality control threshold for head motion (mean framewise displacement (FD) < 0.2 mm) (Supplementary Fig. 22). (*ii*) To mitigate the impact of uneven sample and site distributions across ages, a balanced sampling strategy was employed to ensure uniformity in participant and site numbers (N = 6,770, resampling 1,000 times) (Supplementary Fig. 23). (*iii*) To validate reproducibility of our results, a split half approach was adopted (Supplementary Fig. 24). (i*v*) To examine the potential effects of data samples, a bootstrap resampling analysis was performed (1,000 times, Supplementary Fig. 25). (*v*) To examine the potential effects of specific sites, a leave-one-site-out (LOSO) analysis was conducted (Supplementary Fig. 26). The results of these sensitive analyses were quantitatively assessed in comparison to the main results (Supplementary Tables 7-12).

Specifically, a series of 80 points at one-year intervals was sampled for each curve, and Pearson’s correlation coefficients were then calculated between the corresponding curves (Supplementary Table 7). At both global and system levels, all growth curves in the sensitivity analyses exhibited a high degree of correlations with those shown in the main results (r = 0.97-1.0 for global mean of the connectome; r = 0.98-1.0 for global variance of the connectome; r = 0.99-1.0 for global system segregation; r = 0.98-1.0 for system segregation of VIS, DA, VA, FP, and DM networks; r = 0.91-1.0 for system segregation of SM networks; r = 0.8-1.0 for system segregation of LIM networks, except for r = 0.51 of the balanced resampling analysis; all p_FDR_ < 10^-5^). At the regional level, the lifespan growth axes in the sensitivity analyses were highly spatially associated with that shown in the main results (all r = 0.94-1.0, p_spin_ < 0.0001). The similar results of growth rate are shown in Supplementary Table 8. We observed consistent results when the sampling was obtained with six-month intervals (160 points) and monthly intervals (1,000 points) (Supplementary Tables 9-12).

## Discussion

Using a large multimodal structural and task-free fMRI dataset from 33,250 individuals aged 32 postmenstrual weeks to 80 years, we mapped the growth patterns of the functional connectome across the human lifespan at the global, system, and regional levels. We charted the multiscale, nonlinear growth curves of the functional connectome and revealed previously unidentified key growth milestones. To provide a lifespan characterization of functional brain systems, we created age-specific atlases spanning 32 postmenstrual weeks to 80 years of age to serve as a foundational resource for future research. Using three representative disease datasets of ASD, MDD, and AD, we explored the utility of the connectome-based normative model in capturing individual heterogeneity, identifying disease biotypes, and performing classification and prediction analyses within these clinical populations, highlighting their potential to advance our understanding of neuropsychiatric disorders.

At the global level, we observed continuous nonlinear changes in the global mean and variance of functional connectivity across the life cycle, peaking in the late fourth and late third decades, respectively. Similarly, the growth curve of global brain structure shows a pattern of increase followed by decline, albeit peaking earlier ^18^. Taken together, these functional and anatomical findings suggest that the human brain remains in a state of dynamic adaptation throughout the lifespan. At the systems level, an intriguing observation is that the DM and FP networks, relative to other networks, undergo more rapid development of system segregation during infancy, childhood, and adolescence, peak later, and decline precipitously during aging. The accelerated early development of these networks can be attributed to their initially less organized functional architecture in utero ^53, 83^ and the subsequent need for rapid postnatal development to support the emergence and development of advanced cognitive functions ^8, 84, 85^. Moreover, the increased susceptibility of these networks to accelerated decline during aging may be exacerbated by their increased sensitivity to environmental, genetic, and lifestyle factors, as well as neurodegenerative agents such as amyloid-β and tau ^86–89^. At the regional level, our results validate and extend the replicable findings of Luo and colleagues ^32^, who, using four independent datasets, observed an increase in FCS in primary regions and a decrease in higher-order regions from childhood to adolescence. Furthermore, the life-cycle growth curves of regional FCS are constrained by their positions along the S-A axis, highlighting the role of the S-A axis as a key organizational principle that influences cortical development and aging ^66^.

Emerging evidence increasingly implicates abnormal interregional brain communication and global network dysfunction as critical factors in the pathogenesis of various neuropsychiatric disorders ^13, 15, 16^. After establishing lifespan growth curves, we focused on characterizing the degree to which individual functional metrics deviated from established population norms. This analysis provided preliminary insights into the clinical utility of our connectome-based normative models. Using age- and sex-normalized metrics, we first elucidated individual heterogeneity in functional brain deviations at the global, system, and regional levels across three clinically relevant populations, namely ASD, MDD, and AD. Through subtype analysis based on individual deviation scores, we validated the potential of the connectome-based normative models to parse complex intragroup heterogeneity and enhance the prediction of disease discrimination and clinical symptoms. A biological exploration of the underlying causes of positive and negative deviations in individual functional brain connectomes would provide valuable insights into the similarities and differences between disparate clinical disorders ^78^.

Furthermore, future studies could include more disease cohorts with large sample sizes to allow transdiagnostic comparisons between disorders. It is important to note that considerable work is still needed to effectively translate growth charts and their derived heterogeneity metrics into clinical utility ^18, 90, 91^. Therapeutically, the incorporation of individual functional deviations along with finely stratified subtypes may improve the efficacy of interventions using connectome-guided transcranial magnetic stimulation ^92^. In summary, the integration of the connectomic framework with normative growth curves provides an unprecedented opportunity to study brain network dysfunction in clinical populations.

A promising avenue to explore for future research is the interaction between lifespan growth curves of brain networks under different modalities. This interaction could be investigated by examining how different structural and functional connectivity metrics coevolve across the lifespan and whether there are similar or variable temporal key points within these curves. It would be valuable to determine whether milestones of the structural connectome precede those of the functional connectome, thereby providing an anatomical scaffold for the dynamic maturation of functional communication. Furthermore, identifying the critical physiological factors that shape growth patterns across the lifespan is a complex but essential endeavor. Recent evidence suggests that population-based life-cycle trajectories of cortical thickness align with patterns of molecular and cellular organization, with varying degrees of biological explanation at different life stages ^93^. A genome-wide association meta-analysis by Brouwer et al. ^94^ identified common genetic variants that influence the growth rates in cortical morphology development or atrophy across the lifespan. These findings underscore the necessity of a multifaceted approach encompassing anatomical, genetic, molecular, and metabolic methodologies to elucidate the complex factors that regulate typical and atypical alterations in the human brain connectome.

A number of challenges warrant further consideration. First, the data used to delineate lifespan growth patterns in the current study were aggregated from existing neuroimaging datasets, which are disproportionately derived from European, North American, Asian, and Australian populations. This geographic bias has also been found in other neuroimaging normative references or big data studies, such as those involving cortical morphology growth maps ^18^ and genome-wide association studies of brain structure across the lifespan ^94^. Future research should include more neuroimaging cohort studies designed to achieve a balanced representation of diverse ethnic populations ^95^. In addition, it is critical to consider the diversity of environmental factors, such as socioeconomic status, education level, industrialization, and regional culture, which pose potential challenges to the application of lifespan trajectories. Second, as previously outlined by Bethlehem et al. ^18^, we also encountered challenges related to the uneven age distribution of the neuroimaging sample, particularly with the underrepresentation of the infant and middle-aged (30-40 years) populations. It is evident that functional changes in the uterus are dramatic, however, the paucity of available fetal fMRI data limits our understanding of this critical period. Future research should complement the current models with more neuroimaging data, especially from the fetal stages. Third, the presence of artifacts and low signal-to-noise ratios in fMRI images of the orbitofrontal cortex, partly due to head movement and magnetic field inhomogeneity, represents a significant challenge ^96, 97^. The development of advanced imaging techniques and algorithms will be crucial for addressing this issue. Fourth, adjusting for multisite effects in retrospective data represents another significant challenge. Studies have shown that incorporating site variables as random effects in models, rather than the use of ComBat, is a more effective approach in normative modeling ^18, 98, 99^. Therefore, we adopted a conservative analytical approach by modeling site effects as random effects (for a comparison of results using different methods, see Supplementary Result 4 and Supplementary Fig. 27). Future research may benefit from integrating prospective cohort designs, phantom scans, and scans of traveling subjects. Fifth, due to the ambiguity in interpreting negative functional connectivity, we focused on positive connectivity in our main results. Nonetheless, we also analyzed the normative growth patterns of negative connectivity across the lifespan at global, system, and regional levels (Supplementary Result 5 and Supplementary Fig. 28). Sixth, considering the methodological challenges of surface-based analyses in integrating cortical and subcortical structures, we focused on cortical connectomes in our main results. In light of the significance of subcortical structures, we also presented lifespan growth curves of subcortical connectomes using volume-based analysis (Supplementary Result 6 and Supplementary Fig. 29). Seventh, the data used in this study are cross-sectional, which may result in an underestimation of age-related changes in the functional connectome ^100^. Therefore, integrating more densely collected longitudinal data across all ages is essential to accurately characterize lifespan trajectories. Finally, it is anticipated that the connectome-based growth charts established here will serve as a dynamic resource. As more high-quality, multimodal connectome datasets become available, the lifespan normative growth model will be updated accordingly.

## Methods

### Datasets and participants

To delineate the normative growth of the functional connectome in the human brain, we aggregated the available multisite neuroimaging datasets, each containing both 3T structural and task-free fMRI data. For participants with multiple test-retest scans, only the first session was included. The total number of imaging scans collected was 46,178 with 44,030 participants ranging in age from 32 postmenstrual weeks to 80 years. These scans were obtained from 172 sites in 28 datasets. Participant demographics and imaging scan parameters for each site were presented in Supplementary Table 1 and 2, respectively. Written informed consent was obtained from participants or their legal guardians, and the recruitment procedures were approved by the local ethics committees for each dataset.

### Image quality control process

The implementation of a rigorous and standardized quality control procedure is essential to ensure the authenticity of neuroimaging data, thereby enhancing the credibility of growth curves. Previous research has shown that inadequate quality control of MRI scans can diminish the benefits of large sample sizes in detecting meaningful associations ^101^. In this study, we employed a comprehensive four-step data quality control framework that combined automated assessment approaches and expert manual review to assess both structural and functional images across all 46,178 imaging scans from 44,030 participants (Supplementary Figs 1 and 2). This rigorous framework effectively identified imaging artifacts or errors, thereby ensuring the accuracy and reliability of our neuroimaging data.

#### Step 1: Quality control of the raw images

First, we performed a preliminary quality control to filter out low-quality scans with problematic acquisitions. For several publicly available datasets (dHCP, HCP-Development, HCP-Aging, HCP-Young Adult, and ABCD) that provide information on image quality, we performed initial quality control according to their recommended inclusion criteria. For the BCP dataset, each scan was visually reviewed by two neuroradiologists experienced in pediatric MRI. For the other datasets, we conducted automated quality assessment using the MRI Quality Control (MRIQC) tool ^102^, which extracted non-reference quality metrics for each structural (T1w and T2w) and fMRI image. In each dataset, structural images were excluded if they were marked as outliers (more than 1.5 times the interquartile range (IQR) in the adverse direction) in at least three of the following quality metrics: entropy-focus criterion (EFC), foreground-background energy ratio (FBER), coefficient of joint variation (CJV), contrast-to-noise ratio (CNR), signal-to-noise ratio (SNR), and Dietrich’s SNR (SNRd). Similarly, functional images were excluded if they were marked as outliers in three or more of the following quality metrics: AFNI’s outlier ratio (AOR), AFNI’s quality index (AQI), DVARS_std, DVARS_vstd, SNR, and temporal signal-to-noise ratio (tSNR). This step resulted in the exclusion of 838 structural and 963 functional images.

#### Step 2: Determination of whether to pass the entire processing pipeline

Following the initial quality control step, the images were submitted to the pre- and post-processing pipelines. A detailed description of the latter is provided in the “**Data processing pipeline**” section. Any scan that could not pass the entire data processing pipeline was excluded, resulting in the removal of 2,910 structural and 2,969 functional images.

#### Step 3: Surface quality control and head motion control

For structural images, the Euler number was employed to assess the quality of the reconstructed cortical surface. The Euler number is a mathematical concept that summarizes the topological complexity of a surface and, can be calculated as 2-2n, where n represents the number of defects such as holes or handles. A high Euler number represents a surface with fewer defects, indicating high-quality cortical surface reconstruction. The Euler number is a reliable and quantitative measure that can be used to identify images unsuitable for analysis ^18, 101, 103^. Similarly, the images with an Euler number magnitude less than 1.5 times the IQR in the adverse direction from the study-specific distribution (Q1–1.5*IQR) were identified as outliers and excluded. For functional images, scans with large head motion (mean FD > 0.5 mm, or frames with FD over 0.5 mm > 20%) were excluded, along with scans with fewer than 100 final time points or a ratio of final time points to original time points < 90%. In total, 2,117 structural images and 3,573 functional images were excluded.

#### Step 4: Visual double-check

During the initial three QC steps using automated assessment approaches, 5,865 scans with structural imaging problems and 7,505 scans with functional imaging problems were excluded. To further ensure the quality of the remaining scans, we performed a detailed and comprehensive visual check QC. (1) A visual QC team was assembled, comprising of four anatomically trained experts: Q.W., Q.Y., C.P., and L.S.. For each participant who had passed the automated QC steps, three 2D pictures were generated (one for structural MRI images and two for functional MRI images). (2) Based on these images, L.S. conducted the initial round of visual QC on both structural and functional data for all participants, recording the IDs of those with quality errors. (3) The pictures were then distributed evenly among Q.W., Q.Y., and C.P. for a secondary evaluation. The IDs of the participants exhibiting quality defects were documented. The final list of participants who were excluded was determined based on the combination of these records. Throughout the process, the QC team engaged in in-depth discussions to ensure that the exclusion criteria were consistently applied across members. The exclusion criteria were as follows: The T1-weighted structural images were primarily evaluated for artifacts and quality of cortical segmentation, reconstruction, and registration. For participants with T2-weighted images, those with abnormal myelination distribution (as measured by the T1/T2 ratio) were also excluded. Functional images were assessed for brain coverage, functional-to-structural and functional-to-standard space registration quality, and volume-to-surface mapping quality. Participants were excluded if any of these issues were present. A comprehensive tutorial on visual QC procedures is available at https://github.com/sunlianglong/BrainChart-FC-Lifespan/blob/main/QC/README.md. In this step, 651 structural images and 1,153 functional images were excluded. Finally, only scans that successfully passed QC for both functional and structural images were retained.

The Application of the rigorous criteria outlined above resulted in the exclusion of 10,231 scans in 9,564 participants. The final sample included 33,250 healthy participants (33,250 cross-sectional scans and 1,481 longitudinal scans) and 1,216 patients (1,216 cross-sectional scans; 414 patients with ASD, 622 patients with MDD, and 180 patients with AD) with high-quality functional and structural images.

### Data processing pipeline

#### (i) Structural data preprocessing

Despite our efforts to employ a unified structural preprocessing pipeline across all datasets to mitigate the impact of disparate methodologies, the substantial variations in the structure and function of the human brain across the lifespan present a significant challenge. This was particularly evident in the perinatal and infant periods, where the anatomical characteristics differ markedly from those of adults. For example, in six-month-old infants, the contrast between gray and white matter is extremely subtle, and at approximately six months of age, there is a contrast inversion between gray and white matter. These factors greatly complicate the segmentation of brain tissue during this period ^104, 105^. In the absence of a preprocessing pipeline suitable for all stages of life, it is necessary to find appropriate methods for early developmental datasets while ensuring the uniformity of the pipelines in other datasets.

For individuals aged two years and older, we utilized the publicly available, containerized HCP structural preprocessing pipelines (v4.4.0-rc-MOD-e7a6af9) ^106^, which have been standardized through the QuNex platform (v0.93.2) ^107^. Briefly, this pipeline consists of three stages: (1) The PreFreeSurfer stage focused on the normalization of anatomical MRI data and involved a sequence of preprocessing steps that included brain extraction, denoising, and bias field correction on anatomical T1 weighted (T1w) and T2 weighted (T2w) MRI data (if T2w data were available). (2) The FreeSurfer stage aimed to create cortical surfaces from the normalized anatomical data, including anatomical segmentation; the construction of pial, white, and mid-thickness surfaces; and surface registration to the standard atlas. (3) The PostFreeSurfer stage converted the outputs from the previous steps into the HCP format (CIFTI). The volumes were transformed to the standard MNI space using nonlinear registration, while the surfaces were mapped to the standard fs_LR_32k space using spherical registration and surface downsampling. To mitigate the computational burden of processing the large ABCD dataset, we chose to use the community-shared, preprocessed data released through the ABCD-BIDS Community Collection ^108^ (ABCD collection 3165; https://github.com/ABCD-STUDY/nda-abcd-collection-3165). The multimodal neuroimaging data were preprocessed using the ABCD-HCP pipeline, a variant of the HCP pipeline adapted to better suit the ABCD dataset. Modifications to the ABCD-HCP structural pipeline include volume registration algorithms and bias field correction methods. Further details of these modifications can be found in the online document (https://collection3165.readthedocs.io/en/stable/pipelines/).

For participants in the postmenstrual age range of 32 to 44 weeks from the dHCP study, we applied the officially recommended dHCP structural pipelines ^109^, which have been specifically designed to account for the substantial differences between neonatal and adult MRI data. This HCP-style pipeline included the following steps: (1) bias correction and brain extraction, which were performed on the motion-corrected, reconstructed T2w images; (2) tissue segmentation; (3) cortical reconstruction of the white matter surface; (4) surface topology correction; (5) generation of pial and mid-thickness surfaces; (6) generation of inflated surfaces derived from the white matter surface through an expansion-based smoothing process; and (7) projection of the inflated surface onto a sphere for surface registration. Furthermore, we used the officially recommended iBEAT V2.0 pipelines ^110^ for participants aged from 0-2 years (all from the BCP study). This pipeline, which is optimized for preprocessing early-age neuroimaging data based on advanced algorithms, has shown superior performance in tissue segmentation and cortical reconstruction for BCP datasets compared to alternative approaches ^110^. The main steps of this pipeline included (1) inhomogeneity correction of T1w/T2w images; (2) skull stripping and cerebellum removal (for participants with incomplete cerebellum removal, frame-by-frame manual corrections were performed); (3) tissue segmentation; (4) cortical surface reconstruction; (5) topological correction of the white matter surface; and (6) final reconstruction of the inner and outer cortical surfaces. To ensure consistency in data preprocessing, we employed the iBEAT pipeline for structural image preprocessing of participants aged 2-6 years (53 scans, representing 13% of the total BCP cohort) from the BCP site.

The individual cortical surface obtained from the dHCP and iBEAT V2.0 structural pipelines were aligned with the adult fs_LR_32k standard space using a three-step registration method (Supplementary Fig. 3). For participants aged 32 to 44 postmenstrual weeks, the following steps were implemented. (1) Individual surfaces were registered to their respective postmenstrual week templates ^111^. (2) Templates for 32-39 postmenstrual weeks and 41-44 postmenstrual weeks were registered to the 40-week template. (3) The 40-week template was then registered to the fs_LR_32k surface template. For participants aged 1-24 months, the following steps were undertaken. (1) Individual surfaces were registered to their corresponding monthly age templates ^112^. (2) All monthly templates were registered to the 12-month template. (3) The 12-month template was then registered to the fs_LR_32k surface template. A supplementary analysis was conducted to validate the normative growth pattern of the global functional connectome, which involved avoiding cross-age surface registration (Supplementary Result 7 and Supplementary Fig. 30).

#### (ii) Functional data preprocessing in volumetric space

For individuals aged two years and older, the HCP functional preprocessing pipelines were employed ^106^. The fMRIVolume stage consisted of the following steps. (1) Slice timing correction: This step was applied to single-band acquisitions, as multi-band acquisitions did not require slice timing correction. (2) Motion correction: EPI images were aligned to the single-band reference image using 6 DOF FLIRT registration. In cases where the single-band imaging data were not available, the first frame of the fMRI data was used as the reference. The motion parameters, including translations, rotations, and their derivatives were recorded. Additionally, the demeaned and linearly detrended parameter was provided for nuisance regression analysis. (3) EPI distortion correction: Geometric distortion correction was conducted using either the opposite-phase encoded spinLecho images (when LR-RL or AP-PA encoded acquisitions were available) or the regular (gradient-echo) fieldmap images (when fieldmap acquisitions were available). If neither image was available, this step was skipped. (4) Anatomical registration: The fMRI images were registered to the T1w image using 6 DOF FLIRT with boundary-based registration (BBR). (5) Intensity normalization: The fMRI data, masked by the final brain mask generated by the PostFreeSurfer structural pipeline, were normalized to a 4D whole-brain average of 10,000.

For participants in the postmenstrual age range of 32 to 44 weeks from the dHCP study, we applied the dHCP functional pipelines ^113^. Building on the foundation of the HCP pipeline and the FSL FEAT pipeline, this pipeline was tailored to address the unique challenges associated with neonatal fMRI data. The key components of the pipeline included the following steps. (1) Fieldmap preprocessing, which included estimation of the susceptibility distortion field based on the opposite-phase encoded spinLecho images and subsequent alignment of this field to the functional data. (2) Registration, which included BBR of the fieldmap magnitude to the T2w image, BBR of the single-band reference image to the T2w image with incorporation of field map-based distortion correction, and 6 DOF FLIRT registration of the first volume of the functional multiband EPI to the single-band reference image. (3) Susceptibility and motion correction, which included slice-to-volume motion correction, motion-by-susceptibility distortion correction, and estimation of motion nuisance regressors. These steps resulted in distortion-corrected and motion-corrected 4D multiband EPI images in the T2w native volumetric space. For participants from the BCP cohort, we implemented several steps to obtain preprocessed volumetric fMRI data. (1) Motion correction: functional images were aligned to the single-band reference image using 6 DOF FLIRT registration. In the absence of a single-band reference, the mean functional images (with all frames aligned to the first frame) were employed as the reference. (2) Distortion correction: we performed distortion correction based on the opposite-phase encoding (AP-PA) spinLecho images. This step was only performed for participants with available images. (3) EPI to anatomical registration: the reference image was aligned to the anatomical image (T1w or T2w) using 6 DOF FLIRT registration.

#### (iii) Functional data preprocessing in surface space

In the fMRISurface stage of the HCP functional pipeline, the goal was to project the volume time series onto the standard CIFTI grayordinates space. For the data from the dHCP and BCP cohorts, we followed the same steps of the HCP preprocessing pipeline to achieve an accurate representation of cortical BOLD signals on the surface. Specifically, the fMRI volumetric data in the cortical cortex were separated into left and right hemispheres and mapped onto each participant’s mid-thickness surfaces using a partial-volume weighted, ribbon-constrained volume-to-surface mapping algorithm ^106^. Subsequently, the time courses were then transferred from the individual’s native space to the fs_LR_32k standard space using each participant’s surface registration transformations from the structural preprocessing stage.

#### (iv) Functional data postprocessing

For the ABCD dataset, the ABCD-HCP functional pipeline used DCANBOLDProcessing software (https://collection3165.readthedocs.io/en/stable/pipelines/) to reduce spurious variance that is unlikely to reflect neural activity. For other datasets, the preprocessed fMRI data were post-processed using SPM12 (v6470) and GRETNA (v2.0.0) with a uniform pipeline. Specifically, the following steps were initially conducted on the time series for each vertex in fs_LR_32k space (59,412 vertices in total): linear trend removal, regression of nuisance signals (24 head motion parameters, white matter signal, cerebrospinal fluid signal, and global signal), and temporal bandpass filtering (0.01–0.08 Hz). To mitigate the effects of head motion, the motion censoring was further implemented. This process involved discarding volumes with an FD greater than 0.5 mm and adjacent volumes (one before and two after). To maintain the temporal continuity of the fMRI time series, we subsequently filled these censored frames using a linear interpolation.

Participants with more than 20% of frames exceeding the 0.5 mm FD threshold were excluded from our study. Surface-based smoothing was then applied using a 6-mm full-width at half-maximum (FWHM) kernel. Finally, the data were resampled to a mesh of 2,562 vertices (corresponding to the fsaverage4 standard space) for each hemisphere using the HCP Workbench *metric-resample* command. The removal of the medial wall resulted in a combined total of 4,609 vertices exhibiting BOLD signals on both the left and right hemisphere surfaces.

### Construction of the age-specific and individualized functional atlases across the lifespan

#### (i) Construction of population-level age-specific atlases

To improve the precise mapping of individual-specific functional networks across the lifespan, we first developed a Gaussian-weighted iterative age-specific group atlas (GIAGA) generation approach (Supplementary Fig. 6a) to create a set of age-specific population-level functional atlases (Fig. 2a, Supplementary Figs 7 and 8). Given the dramatic functional changes that occur during early development ^57^, we prioritized the generation of finer age-specific atlases for these stages compared to the later life stages. To this end, we divided all individual scans into 26 different age subgroups, ranging from 32 postmenstrual weeks to 80 years of age. Each age group consisted of cross-sectional data only. Then, we constructed an age-specific functional atlas for each subgroup. A total of 9 atlases were constructed for the perinatal to early infant period, including 4 for perinatal development (34-week, 36-week, 38-week, and 40-week (0-year) atlases) and 5 for the first year of life (1-month, 3-month, 6-month, 9-month, and 12-month atlases). 2 atlases were developed for toddlers (18-month and 24-month atlases), while 9 atlases were created for childhood and adolescence (4-year, 6-year, 8-year, 10-year, 12-year, 14-year, 16-year, 18-year, and 20-year atlases). Finally, 6 atlases were generated for adults and the elderly (30-year, 40-year, 50-year, 60-year, 70-year, and 80-year atlases). A total of 300 participants were randomly selected for each age subgroup. In the event that the available sample size was less than 300, all participants who passed the imaging quality control were included. Further details on the age range, number of participants, and sex ratio for each atlas can be found in Supplementary Table 13.

In recent studies of brain functional organization, Yeo’s 7- and 17-network atlases ^58^ have been widely used to map cortical functional systems ^114^. By including hand sensorimotor areas based on activations in a hand motor task ^115^, Wang and colleagues extended this classical functional parcellation, resulting in an 18-network atlas ^60^. In line with previous studies ^47, 61, 62^, we utilized this updated classic 18-network map as the initial atlas for the construction of age-specific group atlases. The detailed construction process for a given age subgroup (e.g., 17-19 years) was as follows. First, to enrich the dataset for this age subgroup, we included the latter half of the participants from the previous subgroup (15-17 years) and the earlier half of the participants from the subsequent subgroup (19-21 years). We then used the individualized parcellation iteration algorithm proposed by Wang and colleagues ^60^ to map the 18-network atlas to each participant, generating the initial individualized functional parcellations (step 1 in Supplementary Fig. 6a). We then proposed the GIAGA approach. Around the core age (i.e., 18 years) of this given group, we generated a Gaussian probability distribution N(µ, σ^2^) with mean µ = 0 and standard deviation σ = 1 and assigned weights to each participant based on their age position in this Gaussian distribution. The weight quantified the participant’s contribution to the population-level atlas construction, with closer to the core age resulting in a greater contribution. For each vertex, we calculated the cross-participant cumulative probability of belonging to each network and assigned vertex labels to the network with the highest cumulative probability, resulting in an initial age-specific population-level atlas (step 2 in Supplementary Fig. 6a). Finally, steps 1 and 2 were iteratively repeated until the overlap between the current and previous atlases exceeded 95% or the total number of iterations exceeded 10, indicating convergence (step 3 in Supplementary Fig. 6a).

#### (ii) Individualized atlas construction

For a given participant, we used the same iterative parcellation method described above to generate an individualized functional parcellation based on the corresponding population-level atlas specific to the participant’s subgroup (Supplementary Fig. 6b, adapted from ^60^). Briefly, the influence of the population-level atlas on the individual brain varied across participants and across brain regions; therefore, this method made flexible modifications during the construction of the individualized atlas based on the distribution of intersubject variability in the functional connectome and the tSNR of the functional BOLD signals ^60^. Over the iterations, the weight of population-based information was progressively reduced, allowing the final individualized map to be completely driven by the individual-level BOLD data. More information on this iterative functional parcellation approach can be found in the study by Wang and colleagues ^60^.

Notably, given the potential variance of different interindividual variability patterns and tSNR distributions across different age subgroups, we generated an interindividual variability map and a tSNR map for each age subgroup. This was done to improve the accuracy of both the individual and population-level atlases. We divided the time series data of each participant within each age subgroup into two halves. For each half, we computed a vertex-by-vertex functional connectome matrix. This allowed us to obtain the intersubject variability and the intrasubject variability within the subgroup. By regressing the intrasubject variability from the intersubject variability, we obtained a "purified" measure of intersubject variability in the functional connectome ^116, 117^.

#### (iii) Construction of the reference atlas used for comparison

To mitigate the potential bias introduced by specifying a reference atlas for ‘mature age’, we adopted a data-driven approach to construct the reference atlas. Atlas similarity was assessed using the overlap index, which quantifies the proportion of vertices with matching labels between two atlases. For instance, if two atlases have 4,000 vertices with identical labels out of a total of 4,609 vertices, the overlap index would be 4,000/4,609 = 86.8%. We computed the overlap index between each pair of the 26 atlases, resulting in a 26×26 similarity matrix. Hierarchical clustering was applied to this matrix, as shown in Supplementary Fig. 10a. We selected a highly congruent cluster of atlases, including the 18-, 20-, 30-, 40-, 50-, 60-, and 70-year atlases. For each vertex, we assigned the label as the system that had the highest probability of occurrence across these selected atlases, thereby generating the final reference atlas (Supplementary Fig. 10b).

#### (iv) Homogeneity of both the age-specific and personalized functional atlases

We evaluated the functional homogeneity of three parcellation atlases at specific age intervals: the adult-based group atlas established by Yeo et al. ^58^, the age-specific group atlas, and the individual-specific atlas (Supplementary Fig. 9). For each age interval, we performed one-way repeated measures analysis of variance (RANOVA) followed by post hoc multiple comparisons tests to determine whether the homogeneity of the individualized atlas was significantly greater than that of the age-specific group atlas and whether the homogeneity of the age-specific group atlas was significantly greater than that of the adult-based group atlas.

The homogeneity of a system was assessed by calculating the average similarity between every pair of vertices assigned to it. The commonly used metric is within-system homogeneity, which is calculated as the average of Pearson’s correlation coefficients between the time series of all vertex pairs within each system, serving as a measure of internal consistency ^48, 49^. To summarize within-system homogeneity for comparisons across atlases, we averaged the homogeneity values across systems ^49^. For validation, we employed another commonly used metric, the functional profile homogeneity, which defines system similarity as Pearson’s correlation coefficient between the “connectivity profiles” of vertices within a system ^118, 119^. The connectivity profile of a vertex is represented by the connections between this vertex with all other cortical vertices. The global average functional profile homogeneity value was derived by averaging the homogeneity values across all systems ^119^. The RANOVA revealed significant differences in the global average of functional homogeneity across different atlases for any given age interval (all F > 267, p < 10^-25^, Supplementary Fig. 9). Post hoc analysis revealed significant differences in functional homogeneity between every pair of atlases (all p < 10^-8^, individual-specific atlas > age-specific group atlas > adult-based group atlas, Supplementary Fig. 9), regardless of the age groups considered.

### Individualized metrics of the functional connectome at global, system, and regional levels

For each pair of vertices among the 4,609 vertices in the fsaverage4 space, we computed the Pearson’s correlation coefficient to characterize the vertex-by-vertex functional connectivity, resulting in a 4,609×4,609 functional connectome matrix for each participant. All negative functional connectivity strengths were set to zero. For each participant, the global mean of functional connectome was defined as the mean of all 4,609×4,609 connections (edges), and the global variance of functional connectome was defined as the standard deviation of all 4,609×4,609 connections. For validation, we also calculated the global mean of the functional connectome by averaging only the positive-weight edges, which yielded similar lifespan growth patterns (Supplementary Result 8 and Supplementary Fig. 31). At a regional level, the FCS of a given vertex was quantified as the average of the connections with all other vertices.

For a given brain system, an individual’s within-system functional connectivity *FC_W_* was defined as the average connection strength among all vertices within that personalized system. Conversely, the individual’s between-system connectivity *FC_b_* was represented by the average strength of connections between this system a nd all other systems. System segregation ^64^ was determined by calculating the difference between *FC_W_* and *FC_b_*, normalized by *FC_W_*, as described in the following formula:

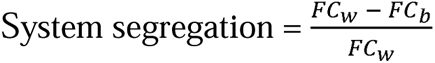

Similarly, global system segregation was defined as the difference between global mean within-system connectivity and global mean between-system connectivity, normalized by global mean within-system connectivity.

The degree of global similarity between an individualized atlas and the reference atlas was quantified by the overlap index. This was defined as the number of vertices with the same label in the two atlases divided by the total number of vertices in both atlases. If there were 4,609 vertices with the same label in two atlases, the overlap index was 4,609/4,609 = 1.0. The degree of similarity between an individualized system and its corresponding system in the reference atlas was quantified using the Dice coefficient.

### Modeling normative growth curves across the lifespan

To estimate the normative growth patterns for various metrics of the functional connectome in healthy individuals combined across cohorts, we applied the GAMLSS ^41, 42^ to the cross-sectional data using the *gamlss* package (version 5.4-3) in R 4.2.0. The GAMLSS procedure were established with two steps: identification of the optimal data distribution, followed by determination of the best-fitting parameters for each functional connectome metric. Using these metric-specific GAMLSS models, we obtained nonlinear normative growth curves and their first derivatives. Furthermore, the sex-stratified growth patterns were revealed. The goodness of fit of the model was confirmed by out-of-sample metrics and visualized by traditional QLQ (quantileLquantile) plots and detrended transformed Owen’s plots. The robustness of the lifespan growth curves was assessed through bootstrap resampling analysis, leave-one-study-out analysis, balanced resampling analysis, and split-half replication analysis.

#### (i) Model data distributions

While the World Health Organization provides guidelines for modeling anthropometric growth charts (such as head circumference, height, and weight) using the Box□Cox t-distribution as a starting point ^41^, we recognized that the growth curves of brain neuroimaging metrics do not necessarily follow the same underlying distributions. For instance, Bethlehem et al. reported that the generalized gamma distribution provided the best fit for brain tissue volumes ^18^. Therefore, we evaluated all continuous distribution families (n=51) for model fitting. To identify the optimal distribution, we fitted GAMLSS with different distributions to four representative global functional metrics (global mean of the connectome, global variance of the connectome, global atlas similarity, and global system segregation) and assessed model convergence. The Bayesian information criterion (BIC) was used to evaluate the fits of the converged models. A lower BIC value indicates a superior fit. As shown in Supplementary Fig. 32, the Johnson’s Su (JSU) distribution consistently demonstrated the optimal fit performance across all the evaluated models.

#### (ii) GAMLSS framework

We performed the GAMLSS procedure with the functional connectome metric as the dependent variable, age as a smoothing term (using the B-spline basis function), sex and in-scanner head motion (HM) as other fixed effects, and scanner sites as random effects. The JSU distribution, which has four parameters: median (*µ*), coefficient of variation (*σ*), skewness (*ν*), and kurtosis (*τ*), was chosen to fit the data distribution. Each functional connectome metric, denoted by *y*, was modeled as:

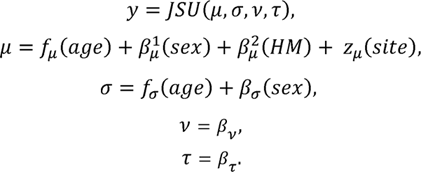

Given the growth complexity across the lifespan, we sought to capture the underlying age-related trends by exploring a range of model specifications. We fitted three GAMLSS models with different degrees of freedom (df = 3-5) for the B-spline basis functions in the location (*µ*) parameters, and set default degrees of freedom (df = 3) for the B-spline basis functions in the scale (*σ*) parameters. Following the practice of previous studies ^18, 80^, only an intercept term was included for the *v* or *τ* parameter. For model estimation, we used the default convergence criterion of log-likelihood = 0.001 between model iterations and set the maximum number of iteration cycles as 200. Finally, the optimal model of a given functional metric was selected based on the lowest BIC value among all converging models. In our study, we did not observe instances of nonconvergence in the GAMLSS models for any metric, including those used in sensitivity analyses.

#### (iii) Goodness of fit of the normative model

To assess the quality of the model fits, we employed a training-test split strategy, which enabled us to recognize the importance of out-of-sample metrics. The dataset was randomly divided into two halves, with one half being used for training (N = 16,663) and the other for testing (N = 16,587). The stratification by site was applied to both halves. Subsequently, the GAMLSS model was refitted using the training set and the model’s goodness of fit was evaluated using the testing set. This procedure was repeated by interchanging the roles of the training and testing sets.

The model’s goodness of fit for the central tendency was assessed using R-squared (R^2^). The calibration of the centiles was evaluated using quantile randomized residuals (also known as randomized z-scores) ^120^. If the modeled distribution closely aligns with the observed distribution, the randomized z-scores should follow a normal distribution, regardless of the shape of the modeled distribution ^121, 122^. We used the ShapiroLWilk test to determine the normality of the distribution of the randomized z-scores, where a *W* value close to 1 indicated good normality.

Additionally, we examined the higher-order moments (skewness and kurtosis) of the randomized residuals to gain deeper insights into the goodness of fit of the normative model ^121^. Skewness values close to 0 indicate symmetrically distributed residuals, showing no left or right bias, and kurtosis values close to 0 indicate a desirable light-tailed distribution. The results demonstrated that nearly all models had skewness and kurtosis values close to 0, with the ShapiroLWilk *W* values consistently above 0.99 (Supplementary Figs 33 and 34, Supplementary Table 14). The R^2^ values for the global connectome mean, global connectome variance, global atlas similarity, and global system segregation were 0.49, 0.48, 0.56, and 0.36, respectively. The R^2^ values for the system segregation of each network ranged from 0.14 to 0.32.

Furthermore, the normalized quantile residuals of the normative model were visually assessed using two diagnostic methods. First, we inspected the plots related to the residuals. As shown in Supplementary Fig. 35, the residuals against the fitted values of *µ* and the index were uniformly distributed around the horizontal line at 0. In addition, the kernel density estimation of the residuals showed an approximately normal distribution, and the normal quantileLquantile (QLQ) plots showed an approximately linear trend with an intercept of 0 and a slope of 1. Second, we used the detrended transformed Owen’s plots of the fitted normalized quantile residuals to evaluate the performance of the models. This function uses Owen’s method to construct a nonparametric confidence interval for the true distribution. As shown in the resulting plots (Supplementary Fig. 36), the zero horizontal line fell within the confidence interval, suggesting that the residuals followed a normal distribution.

#### (iv) Sex differences across the lifespan

In the GAMLSS model, sex was included as a fixed effect to evaluate its impact on the lifespan curves of the functional connectome. We obtained the *µ* and σ coefficients, as well as their standard errors, *T*-values, and *P*-values, for the sex variable using the *summary* function in R as detailed in Supplementary Tables 3 and 4. The estimated *µ* and a coefficients represent the adjusted mean and variance effect of sex on the functional phenotype, considering control variables such as age, head motion (mean FD), and the random effects of scanner site. The *T*-value, calculated as the coefficient divided by its standard error, serves as a statistic to test the null hypothesis that the coefficient is equal to zero (no effect).

### Sensitivity analysis of the connectome-based normative models

The lifespan normative growth patterns were validated at the global, system, and regional levels using various analysis strategies. These analyses addressed key methodological concerns including head motion, the impact of uneven sample and site distributions across ages, replication using independent samples, model stability, and potential effects of specific site. At the global and system level, we quantitatively assessed the similarity between these validated growth patterns and the main results by sampling 80 points at one-year intervals for each growth curve and growth rate and calculating Pearson’s correlation coefficient between the corresponding curves. The sampling was also conducted at six-month intervals (160 points) and monthly intervals (1,000 points). At the regional level, we calculated the spatial association between the lifespan growth axis in the sensitivity analyses and that shown in the main results.

#### (i) Analysis with stricter head motion threshold (mean FD threshold < 0.2 mm)

Previous research has indicated that head motion can significantly impact the quality of brain imaging data ^123–125^. To ensure that our findings were not influenced by the potential effects of head motion, we implemented a stricter quality control threshold, excluding participants with a mean FD exceeding 0.2 mm, and replicated all normative model analyses. Specifically, after excluding 8,756 participants from the initial cohort of 33,250 participants with a 0.5 mm mean FD threshold, we used data from 24,494 participants to validate the lifespan growth curves of the functional brain connectome at the global, system, and regional levels (Supplementary Fig. 22).

#### (ii) Balanced resampling analysis

To address potential biases arising from uneven sample and site distributions across age groups, a balanced sampling strategy was performed (Supplementary Fig. 23). This approach ensured equitable participant and site counts across various age groups through random sampling. Specifically, we divided the entire age range across the lifespan into sixteen age groups (each spanning five years) and then calculated the number of participants and sites for each age group. Besides the age groups under 5 years of age or over 70, the (35, 40] age group had the fewest participants at 464 and the (40, 45] age group contained the fewest sites at 23 (Supplementary Fig. 23-Ia). Thus, we selected all participants from the 23 most populated sites within the (35,40] age group, comprising 457 participants. For other age groups, a random sampling strategy was implemented to include 457 participants from the 23 most populated sites. The resulting distribution of participants and sites across age groups after resampling is shown in Supplementary Fig. 23-Ib.

For global and system metrics, sampling was repeated 1,000 times using the above procedure on a pool of 33,250 participants. For each sampling, we randomly selected 6,770 participants and re-performed the GAMLSS models, resulting in 1,000 sets of growth curves for each metric. We then calculated the 95% CI for these curves, the 95% CI for the peak of the median (50th) centile, and the correlations between the 1,000 median centile lines and the median centile line derived from the original cohort of 33,250 participants. For regional metrics (i.e., FCS), we selected a random resample and recalculated all results, including the normative growth curves and growth rate of the regional FCS, the lifespan growth axis, and the association between the lifespan growth axis and the S-A axis.

#### (iii) Split-half replication analysis

To assess model replicability in independent datasets, a split-half strategy was conducted (Supplementary Fig. 24). Participants were randomly divided into two subgroups, each comprising 50% of the participants (N_Subgroup1_ = 16,663, N_Subgroup2_ = 16,587), with stratification by site. The lifespan normative growth patterns were independently evaluated using Subgroup 1 and Subgroup 2.

#### (iv) Bootstrap resampling analysis

To assess the robustness of the lifespan growth curves and obtain their confidence interval, a bootstrap resampling analysis was performed (Supplementary Fig. 25). This involved the execution of 1,000 bootstrap repetitions using replacement sampling. To ensure that the bootstrap replicates preserved the age and sex proportionality of the original studies, the lifespan (from 32 weeks to 80 years) was segmented into 10 equal intervals and stratified sampling was conducted based on both age and sex. For each functional metric, 1,000 growth curves were fitted and 95% CIs were computed for both the median (50th) centile curve and the inflection points. The 95% CI were calculated based on the mean and standard deviation of the growth curves and growth rates across all repetitions.

#### (v) Leave-one-study-out (LOSO) analysis

To ascertain whether the lifespan growth curves were influenced by specific sites, the LOSO analyses were implemented (Supplementary Fig. 26). In each instance, the samples were removed from one site at a time, the GAMLSS models were refitted and the parameters and growth curves were estimated. We initially compared the curves obtained after excluding the largest site (Site 1 from the UK Biobank dataset, 12,877 participants) with those fitted using the entire dataset (N = 33,250). This reveals that both the growth curves and growth rates were almost identical. The mean and standard deviation across all repetitions were used to calculate the LOSO 95% CIs for both the normative growth curves and growth rates. The narrow CI indicated that our models were robust when data from any single site were removed.

### Clinical relevance of connectome-based normative models in brain disorders

To ascertain the clinical relevance of the established lifespan connectome models, the present study included quality-controlled structural and functional MRI data from three brain disorders. All procedures of quality control, image preprocessing, and network analysis were identical to those used for connectome-based normative modeling. The final analyses comprised data from 591 HCs and 414 patients with ASD from the ABIDE dataset (13 sites), 535 HCs and 622 patients with MDD from the DIDA-MDD dataset (5 sites), and 187 HCs and 180 patients with AD from the MCADI dataset (5 sites).

#### (i) Individual deviation z scores

The standard protocol for normative model ^82^ emphasizes the importance of incorporating some control samples from the same imaging sites as the patients to the testing set. This is done to verify that the observed case□control differences are not due to the analysis with controls in the training set and cases in the testing set ^78, 82^. This approach also allows for the estimation of site effects within the case□control datasets. To establish the normative models for all three disorders using the same set of healthy participants, all the HCs of the three case□control datasets were randomly divided in half (N_train_ = 654; N_test_ = 659). This was done in a stratified manner by age, sex, and site. Lifespan connectome-based normative models were reconstructed by using the training set (N = 32,591), which consisted of half of the HCs (N_train_ = 654) and all samples of other datasets (N = 31,937). The testing set, comprising another half of HCs (N_test_ = 659) and the patient cases, was used as a completely independent set to determine their deviation scores. Specifically, the individual quantile scores were first estimated relative to the normative curves. Subsequently, the deviation z scores were derived using quantile randomized residuals ^120^, an approach that transforms quantiles of the fitted JSU distribution into standard Gaussian distribution z scores. This process was repeated 100 times, generating 100 new models and 100 sets of deviation scores for both the patients and the testing set of the healthy controls. The normality of the distribution of the deviation z scores was assessed and confirmed using a two-tailed KolmogorovLSmirnov test. *P-values* < 0.05 were observed for all functional metrics in all repetitions. Our subsequent analysis was based on these independently derived deviation scores in the HCs (HC_test_) and disease cases.

#### (ii) Stability of deviation scores across 100 repetitions

To quantitatively assess the similarity between the estimated growth curves in 100 distinct normative models and the curves in the main results, we sampled 80 points at one-year intervals for each growth curve and calculated Pearson’s correlation coefficients between the corresponding curves (Supplementary Fig. 15, Supplementary Tables 5 and 6). The curves of all metrics demonstrated a high degree of similarity to the main results (mean r > 0.95, mean MSE < 0.1). To evaluate the stability of individual deviation, we computed the pairwise Pearson’s correlation coefficients and MSEs of the deviation scores among 100 distinct models. The results indicated a high degree of stability in the estimates of individual deviations for patients within specific disease cohorts (mean r > 0.95, mean MSE < 0.2 for all metrics). For case□control group comparison analysis and disease classifications analysis, we replicated the analysis 100 times.

#### (iii) Individual heterogeneity of deviations

Extreme deviations were defined as z > |2.6| (corresponding to a p < 0.005), consistent with the criteria used in previous studies ^75, 76, 78^. The extreme positive and negative deviation scores of each functional metric were calculated for each patient. The percentage map of extreme deviations indicated substantial individual heterogeneity within each disease group (Supplementary Fig. 16).

#### (iv) Disease subtypes identification based on individual functional deviations

Given the substantial individual heterogeneity, we sought to identify subtype differences within each disease cohort (ASD, MDD, and AD) by employing the data-driven k-means clustering algorithm. The deviation features of each patient included the global mean of the connectome, global variance of the connectome, global system segregation, system segregation of each network, and regional level FCS, encompassing a total of 4,619 features. Dimensionality reduction was performed on the normalized features using principal component analysis (PCA). We identified the number of principal components that cumulatively explained more than 95% of the variance, and these components were then used as the features for clustering analysis. The similarity matrix of features across patients was calculated using the Euclidean distance. The optimal number of clusters was determined to be between 2 and 8. A total of 30 different indices were employed from the NbClust package ^126^ to determine the optimal number of clusters. The most frequently identified optimal cluster number was selected as the final cluster count.

#### (v) Case□control difference between patients of each subtype and their matched HCs

The individual deviation scores of patients in each subtype were compared to the median of their matched HCs. For each metric, the significance of the median differences between the patients and HCs was assessed using the MannLWhitney U test. *P*-values were adjusted for multiple comparisons using the BenjaminiLHochberg false discovery rate (FDR) correction across all possible pairwise tests (p < 0.05). For each metric, the case□control difference analysis was repeated 100 times. The proportion of tests that passed the significance threshold in 100 comparisons was reported.

#### (vi) Disease classification based on connectome-based deviations

We performed support vector machines (SVM) analysis to evaluate the ability of connectome-based deviations in discriminating patients from controls. For each disease group, two types of classification models were conducted: classification between all patients and HCs and classification between each subtype of patients and HCs. Each classification model was repeated 100 times. For each time, a 2-fold cross-validation framework was implemented, with each fold alternately serving as the training and test sets. To mitigate the impact of features with greater numeric ranges, we normalized each feature in the training set and applied the resulting parameters to the testing set. We then plotted receiver operating characteristic (ROC) curves and calculated the areas under the curve (AUC) to estimate the classification performance. The statistical significance of the AUC was evaluated using the nonparametric permutation test (1,000 times). During each permutation, the labels of the patients and controls were randomly shuffled before implementing SVM and cross-validation. This process yielded a null distribution of the AUC value, and the *P*-value was computed. Finally, the mean ROC curve was obtained by averaging 100 ROC curves, and the mean AUC value was obtained by averaging 100 AUC values. The codes for the classification analysis were modified from Cui et al. ^127^ (https://github.com/ZaixuCui/Pattern_Classification) and the libsvm software (www.csie.ntu.edu.tw/~cjlin/libsvm/).

#### (vii) Predictions of clinical scores based on connectome-based deviations

Using support vector regression (SVR) with a linear kernel, we sought to assess the ability of the connectome-based functional deviations to predict the clinical scores of patients. A 2-fold cross-validation framework was implemented to estimate the prediction accuracy. For a given disease cohort, the patients were ordered by their target scores and subsequently distributed into alternate folds for training and testing (e.g., 1^st^, 3^rd^, …, to the first fold; 2^nd^, 4^th^, …, to the second fold). Each fold alternately served as the training and test sets. To mitigate the impact of features with greater numeric ranges, we normalized each feature in the training set and applied the resulting parameters to the testing set. The final predictive performance was quantified using Pearson’s correlation coefficients between the predicted and observed clinical scores. The statistical significance of the prediction accuracy was evaluated using the nonparametric permutation test (1,000 times). During each permutation, the observed scores of the patients were randomly shuffled before implementing SVR and cross-validation. This process yielded a null distribution of the correlation coefficients, and the *P*-value was computed. The codes for the prediction analysis were modified from Cui and Gong ^128^ (https://github.com/ZaixuCui/Pattern_Regression_Matlab) and the libsvm software (www.csie.ntu.edu.tw/~cjlin/libsvm/).

## Data availability

The MRI dataset listed in Supplementary Table 1 are partly available at the Adolescent Brain Cognitive Development Study (https://nda.nih.gov/), the Autism Brain Imaging Data Exchange Initiative (https://fcon_1000.projects.nitrc.org/indi/abide/), the Alzheimer’s Disease Neuroimaging Initiative (https://adni.loni.usc.edu/), the Age_ility Project (https://www.nitrc.org/projects/age-ility), the Baby Connectome Project (https://nda.nih.gov/), the Brain Genomics Superstruct Project (https://doi.org/10.7910/DVN/25833), the Calgary Preschool MRI Dataset (https://osf.io/axz5r/), the Cambridge Centre for Ageing and Neuroscience Dataset (https://www.cam-can.org/index.php?content=dataset), the Developing Human Connectome Project (http://www.developingconnectome.org/data-release/second-data-release/), the Human Connectome Project (https://www.humanconnectome.org), the Lifespan Human Connectome Project (https://nda.nih.gov/), the Nathan Kline Institute-Rockland Sample Dataset (https://fcon_1000.projects.nitrc.org/indi/pro/nki.html), the Neuroscience in Psychiatry Network Dataset (https://nspn.org.uk/), the Pediatric Imaging, Neurocognition, and Genetics (PING) Data Repository (http://pingstudy.ucsd.edu/), the Pixar Dataset (https://openfmri.org/dataset/ds000228/), the Strategic Research Program for Brain Sciences (SRPBS) MRI Dataset (https://bicr-resource.atr.jp/srpbsopen/), the Southwest University Adult Lifespan Dataset (http://fcon_1000.projects.nitrc.org/indi/retro/sald.html), the Southwest University Longitudinal Imaging Multimodal Brain Data Repository (http://fcon_1000.projects.nitrc.org/indi/retro/southwestuni_qiu_index.html), and the UK Biobank Brain Imaging Dataset (https://www.ukbiobank.ac.uk/). The dhcpSym surface atlases in aged from 32 to 44 postmenstrual weeks is available at https://brain-development.org/brain-atlases/atlases-from-the-dhcp-project/cortical-surface-template/. The UNC 4D infant cortical surface atlases are available at https://bbm.web.unc.edu/tools/. The fs_LR_32k surface atlas is available at https://balsa.wustl.edu/. The subcortical atlases are available at https://github.com/yetianmed/subcortex. The brain charts and lifespan developmental atlases are shared online via GitHub (https://github.com/sunlianglong/BrainChart-FC-Lifespan).

## Code availability

The codes for this manuscript are available on GitHub (https://github.com/sunlianglong/BrainChart-FC-Lifespan). Software packages used in this manuscript include MRIQC v0.15.0 (https://github.com/nipreps/mriqc), QuNex v0.93.2 (https://gitlab.qunex.yale.edu/), HCP pipeline v4.4.0-rc-MOD-e7a6af9 (https://github.com/Washington-University/HCPpipelines/releases), ABCD-HCP pipeline v1 (https://github.com/DCAN-Labs/abcd-hcp-pipeline), dHCP structural pipeline v1 (https://github.com/BioMedIA/dhcp-structural-pipeline), dHCP functional pipeline v1 (https://git.fmrib.ox.ac.uk/seanf/dhcp-neonatal-fmri-pipeline), iBEAT pipeline v1.0.0 (https://github.com/iBEAT-V2/iBEAT-V2.0-Docker), MSM v3.0 (https://github.com/ecr05/MSM_HOCR), FreeSurfer v6.0.0 (https://surfer.nmr.mgh.harvard.edu/), FSL v6.0.5 (https://fsl.fmrib.ox.ac.uk/fsl/fslwiki), Connectome Workbench v1.5.0 (https://www.humanconnectome.org/software/connectome-workbench), MATLAB R2018b (https://www.mathworks.com/products/matlab.html), SPM12 toolbox v6470 (https://www.fil.ion.ucl.ac.uk/spm/software/spm12), GRETNA toolbox v2.0.0 (https://www.nitrc.org/projects/gretna), BrainNet Viewer toolbox v 20191031 (https://www.nitrc.org/projects/bnv), cifti-matlab toolbox v2 (https://github.com/Washington-University/cifti-matlab), HFR_ai toolbox v1.0-beta-20181108 (https://github.com/MeilingAva/Homologous-Functional-Regions), System segregation code (https://github.com/mychan24/system-segregation-and-graph-tools), Python v3.8.3 (https://www.python.org), neuroharmonize package v2.1.0 (https://github.com/rpomponio/neuroHarmonize), scikit-learn package v1.1.3 (https://scikit-learn.org). R v4.2.0 (https://www.r-project.org), GAMLSS package v5.4-3 (https://www.gamlss.com/),and ggplot2 package v3.4.2 (https://ggplot2.tidyverse.org/).

## Acknowledgments

This work was supported by grants from the National Natural Science Foundation of China (82021004, 82327807, and 31830034 to Y.HE.), the scientific and technological innovation 2030 - the major project of the Brain Science and Brain-Inspired Intelligence Technology (2021ZD0200500 to Q.D. and 2022ZD0211500 to M.R.X.), the Changjiang Scholar Professorship Award (T2015027 to Y.HE.), the Beijing Natural Science Foundation (JQ23033 to M.R.X.), the Beijing United Imaging Research Institute of Intelligent Imaging Foundation (CRIBJZD202102 to M.R.X.), the National Natural Science Foundation of China (31521063 and 31221003 to Q.D., 82071998 to M.R.X., T2325006 to G.L.G., 82202245 to Q.L.L., 81971690 to X.H.L., 32130045 to S.Z.Q., 81571062 and 82172018 to Y.L., 81471120 to X.Z., 61633018 to Y.HAN., 81901101 to P.W., 81400890 to D.W.W., 81920108019, 82330058, 91649117, 81771344, and 81471251 to S.J.Q.), the Beijing Brain Initiative of Beijing Municipal Science & Technology Commission (Z181100001518003 to S.T.), the Fund of Shenzhen Institute for Neuroscience Research (S.T.), the Science and Technology Plan Project of Guangzhou (2018-1002-SF-0442 to S.J.Q.), the Guangzhou Key Laboratory (09002344 to S.J.Q.), and the Key R&D Program of Sichuan Province (2023YFS0076 to T.L.C.). We are grateful to the Adolescent Brain Cognitive Development (ABCD) Study, the Autism Brain Imaging Data Exchange (ABIDE) Initiative, the Alzheimer’s Disease Neuroimaging Initiative (ADNI), the Age_ility Project, the Baby Connectome Project (BCP), the Brain Genomics Superstruct Project (BGSP), the Calgary Preschool MRI Dataset, the Cambridge Centre for Ageing and Neuroscience (Cam-CAN) Dataset, the developing Human Connectome Project (dHCP), the Human Connectome Project (HCP), the Lifespan Human Connectome Project (HCPA & HCPD), the Nathan Kline Institute-Rockland Sample (NKI-RS) Dataset, the Neuroscience in Psychiatry Network (NSPN) Dataset, the Pixar Dataset, the Southwest University Adult Lifespan Dataset (SALD), the Southwest University Longitudinal Imaging Multimodal (SLIM) Brain Data Repository, the UK Biobank (UKB) Brain Imaging Dataset, the Disease Imaging Data Archiving: major depressive disorder (DIDA-MDD) Working Group (PI: Yong He, Lingjiang Li, Jingliang Cheng, Qiyong Gong, Ching-Po Lin, Jiang Qiu, Shijun Qiu, Tianmei Si, Yanqing Tang, Fei Wang, Peng Xie, Xiufeng Xu, and Mingrui Xia), the Multi-center Alzheimer Disease Imaging (MCADI) Consortium (PI: Yong Liu, Xi Zhang, Yuying Zhou, Ying Han, and Qing Wang). We thank the National Center for Protein Sciences at Peking University in Beijing, China, for assistance with MRI data acquisition.

ABCD: data used in the preparation of this article were obtained from the Adolescent Brain Cognitive Development (ABCD) Study (https://abcdstudy.org), held in the NIMH Data Archive (NDA). This is a multisite, longitudinal study designed to recruit more than 10,000 children age 9-10 and follow them over 10 years into early adulthood. The ABCD Study® is supported by the National Institutes of Health and additional federal partners under award numbers U01DA041048, U01DA050989, U01DA051016, U01DA041022, U01DA051018, U01DA051037, U01DA050987, U01DA041174, U01DA041106, U01DA041117, U01DA041028, U01DA041134, U01DA050988, U01DA051039, U01DA041156, U01DA041025, U01DA041120, U01DA051038, U01DA041148, U01DA041093, U01DA041089, U24DA041123, U24DA041147. A full list of supporters is available at https://abcdstudy.org/federal-partners.html. A listing of participating sites and a complete listing of the study investigators can be found at https://abcdstudy.org/consortium_members/. ABCD consortium investigators designed and implemented the study and/or provided data but did not necessarily participate in the analysis or writing of this report. This manuscript reflects the views of the authors and may not reflect the opinions or views of the NIH or ABCD consortium investigators. The ABCD data repository grows and changes over time. The ABCD data used in this report came from https://nda.nih.gov/edit_collection.html?id=3165, shared by the DCAN Labs ABCD-BIDS Community Collection (ABCC) (Collection Investigators: Damien Fair).

ABIDE I: primary support for the work by Adriana Di Martino was provided by the (NIMH K23MH087770) and the Leon Levy Foundation. Primary support for the work by Michael P. Milham and the INDI team was provided by gifts from Joseph P. Healy and the Stavros Niarchos Foundation to the Child Mind Institute, as well as by an NIMH award to MPM ( NIMH R03MH096321).

ABIDE II: primary support for the work by Adriana Di Martino and her team was provided by the National Institute of Mental Health (NIMH 5R21MH107045). Primary support for the work by Michael P. Milham and his team provided by the National Institute of Mental Health (NIMH 5R21MH107045); Nathan S. Kline Institute of Psychiatric Research). Additional Support was provided by gifts from Joseph P. Healey, Phyllis Green and Randolph Cowen to the Child Mind Institute.

ADNI: data used in preparing this article were obtained from the Alzheimer’s Disease Neuroimaging Initiative (ADNI) database (adni.loni.usc.edu). As such, many investigators within the ADNI contributed to the design and implementation of ADNI and/or provided data but did not participate in analysis or writing of this report. A complete listing of ADNI investigators may be found at http://adni.loni.usc.edu/wp-content/uploads/how_to_apply/ADNI_Acknowledgement_List.pdf. Data collection and sharing for this project was funded by the Alzheimer’s Disease Neuroimaging Initiative (ADNI) (National Institutes of Health Grant U01 AG024904) and DOD ADNI (Department of Defense award number W81XWH-12-2-0012). ADNI is funded by the National Institute on Aging, the National Institute of Biomedical Imaging and Bioengineering, and through generous contributions from the following: AbbVie, Alzheimer’s Association; Alzheimer’s Drug Discovery Foundation; Araclon Biotech; BioClinica, Inc.; Biogen; Bristol-Myers Squibb Company; CereSpir, Inc.; Cogstate; Eisai Inc.; Elan Pharmaceuticals, Inc.; Eli Lilly and Company; EuroImmun; F. Hoffmann-La Roche Ltd and its affiliated company Genentech, Inc.; Fujirebio; GE Healthcare; IXICO Ltd.;Janssen Alzheimer Immunotherapy Research & Development, LLC.; Johnson & Johnson Pharmaceutical Research & Development LLC.; Lumosity; Lundbeck; Merck & Co., Inc.; Meso Scale Diagnostics, LLC.; NeuroRx Research; Neurotrack Technologies; Novartis Pharmaceuticals Corporation; Pfizer Inc.; Piramal Imaging; Servier; Takeda Pharmaceutical Company; and Transition Therapeutics. The Canadian Institutes of Health Research is providing funds to support ADNI clinical sites in Canada. Private sector contributions are facilitated by the Foundation for the National Institutes of Health (www.fnih.org). The grantee organization is the Northern California Institute for Research and Education, and the study is coordinated by the Alzheimer’s Therapeutic Research Institute at the University of Southern California. ADNI data are disseminated by the Laboratory for Neuro Imaging at the University of Southern California.

BCP: data used herein is supported by NIH grant (1U01MH110274) and the efforts of the UNC/UMN Baby Connectome Project Consortium. dHCP: data were provided by the developing Human Connectome Project, KCL-Imperial-Oxford Consortium funded by the European Research Council under the European Union Seventh Framework Programme (FP/2007-2013) / ERC Grant Agreement no. [319456]. We are grateful to the families who generously supported this trial.

HCP: data were provided by the Human Connectome Project, WU-Minn Consortium (Principal Investigators: David Van Essen and Kamil Ugurbil; 1U54MH091657) funded by the 16 NIH Institutes and Centers which support the NIH Blueprint for Neuroscience Research; and by the Mc-Donnell Center for Systems Neuroscience at Washington University.

HCP Lifespan: data used in this publication was supported by the National Institute of Mental Health of the National Institutes of Health under Award Number U01MH109589 and by funds provided by the McDonnell Center for Systems Neuroscience at Washington University in St. Louis. The content is solely the responsibility of the authors and does not necessarily represent the official views of the National Institutes of Health.

NKI-RS: Funding for key personnel provided in part by the New York State Office of Mental Health and Research Foundation for Mental Hygiene. Additional project support provided by the NKI Center for Advanced Brain Imaging (CABI), the Brain Research Foundation (Chicago, IL), the Stavros Niarchos Foundation, and NIH grant P50 MH086385-S1.

NSPN: The NSPN study was funded by a Wellcome Trust award to the University of Cambridge and the University College London.

UK Biobank: this research has been conducted using data from UK Biobank (www.ukbiobank.ac.uk). UK Biobank is generously supported by its founding funders the Wellcome Trust and UK Medical Research Council, as well as the Department of Health, Scottish Government, the Northwest Regional Development Agency, British Heart Foundation, and Cancer Research UK.

## Competing interests

The authors declare that they have no competing interests.

## Author contributions

L.L.S. and Y.H. conceptualized the study. Y.H. supervised the project. L.L.S., T.D.Z., X.Y.L., M.R.X., and Y.H. designed the methodology. L.L.S. developed visualizations. Q.L.L., X.H.L., D.N.D., Z.L.Z., Z.L.X., and Z.X.C. provided guidance on data analysis and result interpretation. L.L.S., X.Y.L., Q.W., C.X.P., Q.Y., Q.L.L., Y.H.X. and M.R.X. performed data quality control; G.L.G., Y.C.B., P.D.C., R.C., Y.C., T.L.C., J.L.C., Y.Q.C., Z.J.D., Y.D., Y.Y.D., Q.D., J.H.G., Q.Y.G., Y.H., Z.Z.H., C.C.H., R.W.H., L.J.L., C.P.L., Q.X.L., B.S.L., C.L., N.Y.L., Y.L., J.L., L.L.M., W.W.M., S.Z.Q., J.Q., T.M.S., S.P.T., Y.Q.T., S.T., D.W.W., F.W., J.L.W., P.W., X.Q.W., Y.P.W., D.T.W., Y.K.W., P.X., X.F.X., L.Y.Y., H.B.Z., X.Z., G.Z., Y.T.Z., S.Y.Z collected a subset of the data for this study. L.L.S. and Y.H. wrote the manuscript. All authors reviewed the final manuscript.

## Notes

### Competing Interest Statement

The authors have declared no competing interest.

### Summary of Updates

1. Updating samples and sites. 2. Updating disease-related analysis. 3. Updating normative modeling. 4. Conducting a series of sensitivity analyses to validate our main results.

